# An Evolutionarily Conserved Receptor-like Kinases Signaling Module Controls Cell Wall Integrity During Tip-Growth

**DOI:** 10.1101/704601

**Authors:** Jens Westermann, Susanna Streubel, Christina Maria Franck, Roswitha Lentz, Liam Dolan, Aurélien Boisson-Dernier

**Affiliations:** University of Cologne, Biocenter, 50674 Cologne, Germany; Department of Plant Sciences, University of Oxford, South Parks Road, Oxford OX1 3RB, United Kingdom; Department of Plant and Microbial Biology and Zurich-Basel Plant Science Centre, University of Zurich, 8008 Zurich, Switzerland

## Abstract

Rooting cells and pollen tubes – key adaptative innovations that evolved during the colonization and subsequent radiation of plants on land – expand by tip-growth. Tip-growth relies on a tight coordination between the protoplast growth and the synthesis/remodeling of the external cell wall. In root hairs and pollen tubes of the seed plant *Arabidopsis thaliana,* cell wall integrity (CWI) mechanisms monitor this coordination through the Malectin-like receptor kinases (MLRs) such as *At*ANXUR1 and *At*FERONIA that act upstream of the *At*MARIS PTI1-like kinase. Here, we show that rhizoid growth in the early diverging plant, *Marchantia polymorpha*, is also controlled by an MLR and PTI1-like signaling module. Rhizoids, root hairs and pollen tubes respond similarly to disruption of MLR and PTI1-like encoding genes. Thus, the MLR/PTI1-like signaling module that controls CWI during tip-growth is conserved between *M. polymorpha* and *A. thaliana* suggesting it was active in the common ancestor of land plants.

## Introduction

When plants first colonized terrestrial habitats more than 470 million years ago, they had to cope with a relatively different, water-deprived environment, that represented a major driving force for a series of key adaptations^1–3^. Among these, the emergence of rooting cells, such as rhizoids and root hairs^4^, that facilitate substrate anchorage and water/nutrient uptake, and the subsequent evolution of pollen tubes allowing for an efficient water-independent sperm transport for fertilization in seed plants^5^ use tip-growth for rapid expansion. Tip-growth is an extreme form of polar growth and as such, it relies on a tight coordination in both, time and space, between the turgor pressure-driven deformation/loosening of the preexisting cell wall and the secretion of new plasma membrane and cell wall material to allow for expansion^6^. It has become apparent that plant cells have developed cell wall integrity (CWI) maintenance mechanisms to control this coordination and avoid growth cessation or rupture. Research on tip-growing cells of the seed plant model *Arabidopsis thaliana* has revealed CWI signaling pathways that control pollen tube and root hair growth (reviewed in ^7–11)^. CWI maintenance during pollen tube growth is controlled by two pairs of functionally redundant Malectin-like receptor kinases (MLRs) named *At*ANXUR1/2 (*At*ANX1/2)^12, 13^ and *At*BUDDHA’S PAPER SEAL1/2 (*At*BUPS1/2)^14^. These pollen-expressed MLRs can interact with each other and are thought to form a receptor complex for the peptide ligands *At*RAPID ALKALINIZATION FACTOR 4 (*At*RALF4) and *At*RALF19 ^14, 15^. Downstream of these MLRs, CWI is regulated by reactive oxygen species (ROS)-producing NADPH-oxidases *At*REACTIVE-BURST OXIDASE HOMOLOG H/J (*At*RBOHH/J)^16^, the type-one protein phosphatases *At*ATUNIS1/2 (*At*AUN1/2)^17^, as well as the receptor-like cytoplasmic kinase (RLCK) *At*MARIS (*At*MRI)^18^. Apart from the negative regulators *At*AUN1/2, all the above-mentioned proteins positively regulate CWI maintenance. Consistently, their corresponding loss-of-function and knockdown mutants display precocious pollen tube bursting leading to low or no transmission of the mutant alleles by the haploid pollen to the next generation and thus to male sterility. CWI signaling also occurs in Arabidopsis root hairs, where it is governed by the closest homolog of *At*ANX1/2, the MLR *At*FERONIA (*At*FER); *Atfer* mutants develop root hairs that spontaneously burst during growth^19^. Intriguingly, *At*MRI has been shown to also function during root hair CWI maintenance downstream of *At*FER^18^. Concordantly, *AtMRI* is strongly expressed in both pollen and root tissues and *At*MRI protein is located though the plasma membrane of both pollen tubes and root hairs where it is required to maintain CWI^18, 20^. *At*MRI belongs to the Arabidopsis RLCK-VIII subfamily that shares homology with the tomato Pto-interacting protein 1 (PTI1) involved in the Pto-mediated hypersensitive response^21^. Apart from the bursting root hair and pollen tube phenotypes of *Atmri*, and abscisic acid (ABA)-insensitivity of *Atcark1* mutants^22^, no phenotype and therefore no function have been reported for any of the other 9 Arabidopsis PTI1-like RLCK proteins.

Recently, a T-DNA insertional mutant screen for defective rhizoid growth phenotypes in the early-diverging land plant *Marchantia polymorpha* identified mutant alleles of the two unique Marchantia MLR and PTI1-like kinases, *Mp*FERONIA (*Mp*FER) and *Mp*MARIS (*Mp*MRI), respectively^23, 24^. Here, by focusing on *At*MRI and *Mp*MRI and using trans-species complementation assays, we demonstrate that the rhizoid-derived *Mp*MRI and the root hair- and pollen tube-derived *At*MRI can function in the CWI pathways of all three tip-growing cell types. Despite their different function, origin and growth environment, we show that all three cell types respond similarly to disruption and overexpression of the PTI1-like encoding genes. Finally, we show that *Mp*MRI functions downstream of *Mp*FER in the rhizoid CWI pathway, revealing that the CWI control of tip-growth in bryophytes and seed plants relies on a common MLR/PTI1-like receptor-like kinases module that has been conserved during land plant evolution.

## Results

### *Mp*MRI is required for CWI during rhizoid growth

We previously showed that the Arabidopsis PTI1-like *At*MRI controls CWI during tip-growth of root hairs and pollen tubes^18^. A large-scale mutant screen in Marchantia for defects in rhizoid growth revealed that a T-DNA insertion in the genetic locus *Mapoly0051s0094.1*, renamed *MpMRI* (see below), coding for a PTI1-like homolog, may cause a morphologically related loss of CWI phenotype in tip-growing rhizoids^23^. We first reanalyzed the *Mpmri-1* (formerly named *Mppti-1* or ST13.3^23^) mutant phenotype in young gemmalings and confirmed that the mutant develops significantly shorter rhizoids than the wild-type accessions (Fig. 1a,b). *Mpmri-1* also developed fewer intact rhizoids per thallus than Tak-2 (Fig. 2c), because rhizoids frequently burst at their tip, as revealed by brownish spots on the ventral side of thalli grown on solid medium (Fig. 1a, 2d) or cytoplasmic content released by collapsed rhizoids grown in liquid condition (Fig. 1c). To test whether the T-DNA insertion in the *MpMRI* locus (Supplementary Fig. 1) causes the rhizoid bursting phenotype, we transformed the *Mpmri-1* mutant with the native *MpMRI* coding sequence fused to the red fluorescent protein (RFP) under control of the ubiquitously active promoter of Elongation Factor 1 α (*proMp*EF1α). Rhizoids were longer in transformed lines than in mutants (Fig. 2a and Supplementary Fig. 2). Bursting of rhizoids and the resulting brown staining observed in *Mpmri-1* rhizoids was not observed in any of the transformed lines (Fig. 2d and Supplementary Fig. 2). Thus, disruption of the *MpMRI* locus indeed causes defective rhizoid growth in *Mpmri-1* mutants. The functional *Mp*MRI-RFP protein fusion mainly localizes to the plasma membrane of *Mpmri-1* rescued rhizoids, with a weaker fluorescence signal detected in the cytoplasm (Fig. 2e and Supplementary Video 1). This localization pattern resembles the localization of *At*MRI in Arabidopsis tip-growing cells^18, 20^. Therefore, a plasma membrane-localized *Mp*MRI protein is required for CWI in *M. polymorpha* rhizoids.

**Figure 1:**
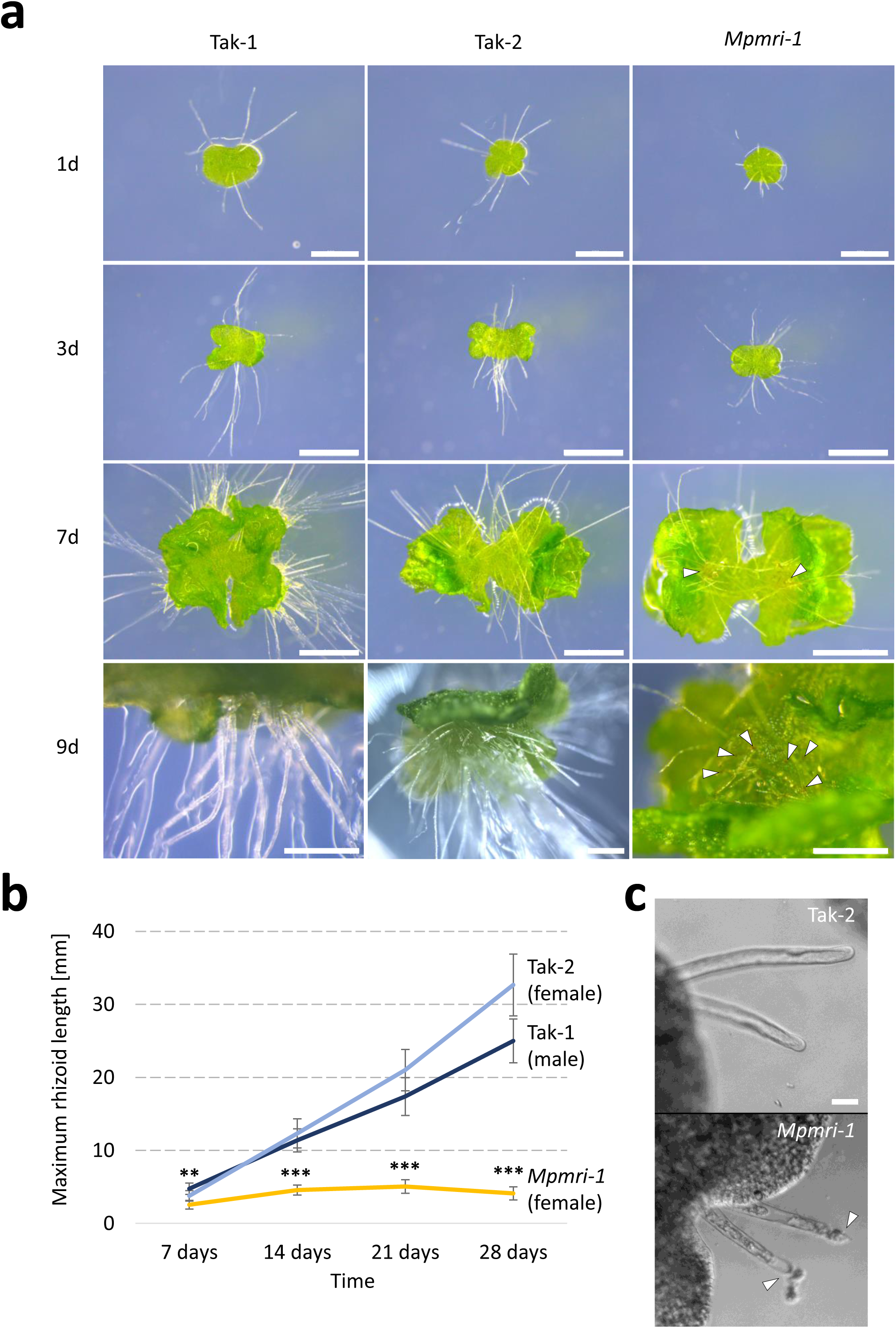
*Mpmri-1* thalli develop short, bursting rhizoids. **(a)** Representative pictures of a time course experiment for thallus and rhizoid development of *Mpmri-1* (female) compared to the Tak-1 (male) and Tak-2 (female) wild-type accessions at t = 1d (scale bar: 500 μm), 3d (scale bar: 1mm), 7d (scale bar: 1mm) and 9d (scale bar: 500 μm) of cultivation. Arrowheads mark the emergence of brownish tissue on the ventral thallus, indicating loss of CWI in emerging rhizoids (observed in 19 out of 20 *Mpmri-1* gemmalings). **(b)** Maximal rhizoid length of Tak-1, Tak-2 and *Mpmri-1* after 7, 14, 21 and 28 days of cultivation (n ≥ 8 thalli/genotype). Significance of *Mpmri-1* dataset against both, Tak-1/2 was tested with a two-tailed, unpaired student’s T-test (**: p <0.01, ***: p<0.001). **(c)** Representative bright field images of rhizoids emerging from 1 d old gemmae. Arrowheads indicate loss of cell wall integrity and release of cytoplasmic content in *Mpmri-1* rhizoids (bottom), as compared to Tak-2 wild-type rhizoids (top). Scale bar = 50 µm.

**Figure 2:**
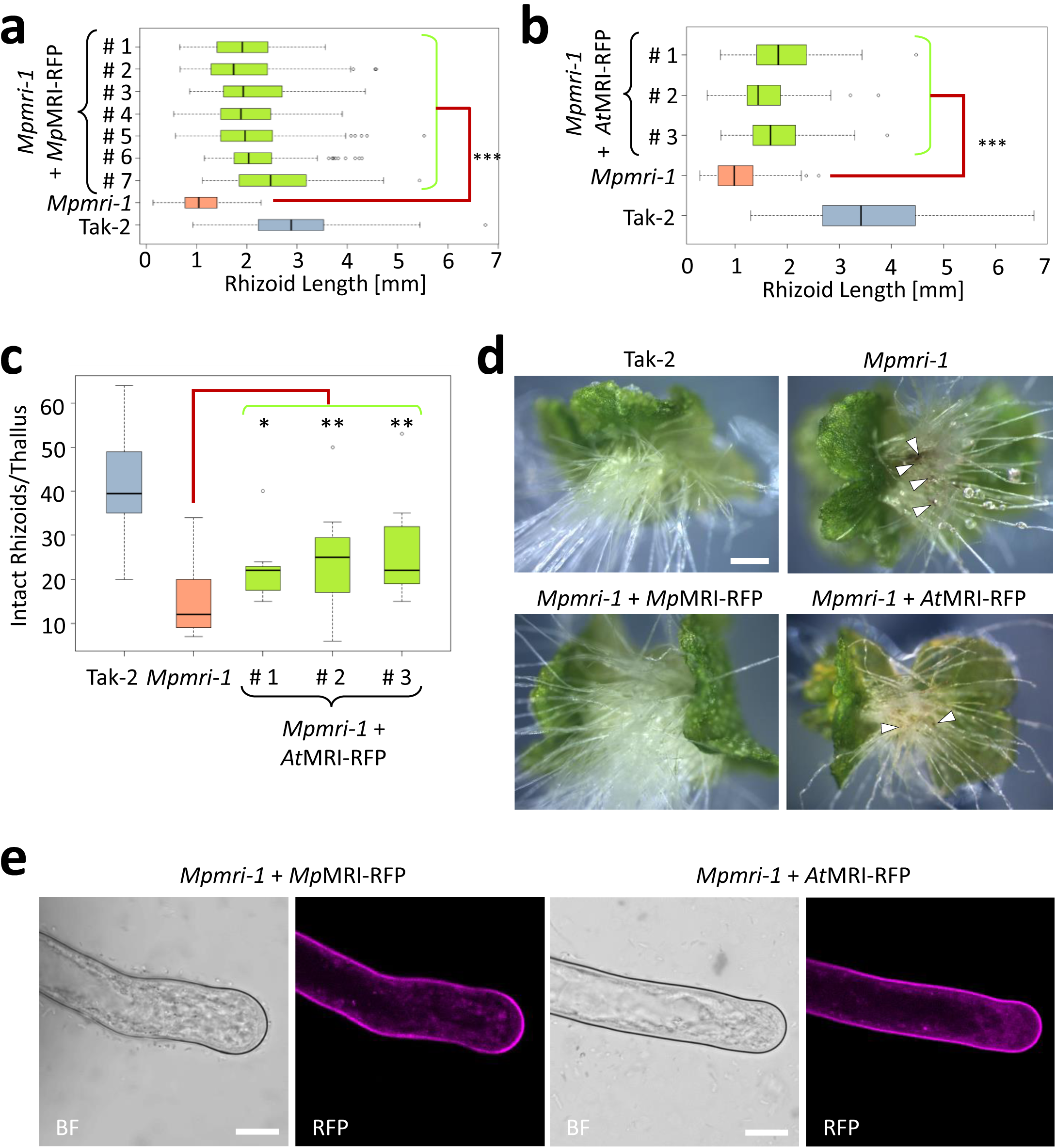
Both, *Mp*MRI and *At*MRI can maintain CWI in growing *Mpmri-1* rhizoids. **(a)** Rhizoid length of 7 d old *Mpmri-1* thalli expressing *proMp*EF1α:*Mp*MRI-RFP (n = 7 independent lines) compared to Tak-2 and non-transformed *Mpmri-1* (n > 10 thalli/line). Significance of all 7 expression lines against non-transformed *Mpmri-1* was tested with a two-tailed, unpaired student’s T-test (***: p<0.001; see also Supplementary Fig. 2). **(b)** Rhizoid length of 7 d old *Mpmri-1* thalli expressing *proMp*EF1α:*At*MRI-RFP (n = 3 independent lines) compared to Tak-2 and non-transformed *Mpmri-1* (n > 10 thalli/independent line). Significance of all 3 lines against non-transformed *Mpmri-1* was tested with a two-tailed, unpaired student’s T-test (***: p<0.001). **(c)** Number of rhizoids of 7 d old *Mpmri-1* thalli expressing *proMp*EF1α:*At*MRI-RFP (n = 3 independent lines) compared to Tak-2 and non-transformed *Mpmri-1* (n ≥ 10 individuals/genotype). Significance of all 3 lines against non-transformed *Mpmri-1* was tested with a two-tailed, unpaired student’s T-test (*: p < 0.05, **: p < 0.01). **(d)** Representative images of the lines analyzed in (a) - (c). Arrowheads depict the frequent loss of CWI at the tip of rhizoids resulting in brownish cytoplasmic material. Scale bar: 500 μm. **(e)** Protein localization of *Mp*MRI-RFP and *At*MRI-RFP in *Mpmri-1* rhizoids, respectively. Bright-field (BF) and RFP filters are indicated. Scale bar: 20 μm. See also Supplementary Video 1,2.

### *At*MRI can maintain CWI during rhizoid growth in Marchantia

Since both *At*MRI and the Marchantia PTI1-like homolog *Mp*MRI control CWI in different tip-growing cells, we tested if their function in CWI was conserved. The sequences of the *At*MRI and *Mp*MRI proteins are 64 % identical (Supplementary Table 1). We first introduced *At*MRI-RFP into the *Mpmri-1* mutant. Expression of *At*MRI-RFP led to a partial, but significant increase in rhizoid length (Fig. 2b) and apparent cell wall integrity (Fig. 2d). The number of intact rhizoids per thallus was significantly higher in all rescue lines as compared to *Mpmri-1* (Fig. 2c). Furthermore, *At*MRI-RFP localized to the plasma membrane of rhizoids just like *Mp*MRI-RFP (Fig. 2e and Supplementary Video 2). Taken together, these results demonstrate that *At*MRI can partially substitute for *Mp*MRI to maintain CWI in *M. polymorpha* rhizoids, despite the independent evolution of both genes for approximately 450-470 million years.

### *Mp*MRI can maintain CWI during pollen tube and root hair growth in Arabidopsis

Next, we tested the hypothesis that the function of PTI1-like proteins has been universally conserved throughout evolution of rooting cells and/or upon acquisition of new functions (*i.e.* between rooting cells and pollen tubes). We first introduced *Mp*MRI fused to the yellow fluorescent protein (YFP) into the heterozygous *Atmri-1*/*At*MRI (herein after referred to as *mri-1/MRI*) background under control of the native *At*MRI promoter (*proAt*MRI), which drives expression in both, root hairs and pollen tubes^17, 18^. Normally, *mri-1* homozygous individuals are never isolated in the self-progeny of heterozygous *mri-1/MRI* plants because *mri-1*-dependent pollen tube bursting prevents transmission of the male *mri-1* allele and thus, production of homozygous *mri-1*/*mri-1* individuals (Supplementary Table 2)^18^. By contrast, 7 homozygous *mri-1*/*mri-1* T2 individuals were isolated out of 200 progenies originating from three independent *mri-1*/MRI T1 lines hemizygous for *Mp*MRI-YFP suggesting that *Mp*MRI expression partially rescues the *mri-1* male gametophytic defect.

To confirm that the partial rescue of *mri-1* male transmission efficiency resulted from the rescue of CWI during pollen tube growth by *Mp*MRI-YFP expression, we performed *in vitro* pollen germination assays on homozygous T3 plants (*mri-1*/*mri-1*; *Mp*MRI-YFP) originating from three independent T1 lines. Pollen bursting rate was lower in all lines expressing *Mp*MRI-YFP and not significantly different from the bursting rates of either wild type, or *mri-1/mri-1* complemented with an *At*MRI-YFP construct (Fig. 3a)^18^. However, the expression of *Mp*MRI-YFP frequently led to abnormal pollen tube morphology *in vitro*, as indicated by swollen apices and branching (n=56 out of 60; Fig. 3b). These abnormalities were not observed in *mri-1* pollen tubes expressing *At*MRI-YFP (Fig. 3b) and may explain the incomplete rescue of male *mri-1-* transmission efficiency by *Mp*MRI-YFP expression. *Mp*MRI-YFP normally localized to the plasma membrane of the pollen tube (Fig. 3b). To exclude that the observed rescue of CWI is not an effect of the artificial *in vitro* conditions, we also determined the seed set of T3 lines. For all T3 lines, there was a significant increase of seed set (Fig. 3c,d). Taken together, these results demonstrate that *Mp*MRI can partially substitute for *At*MRI in *mri-1* pollen tubes despite the different functions and growth environment of rhizoids and pollen tubes.

**Figure 3:**
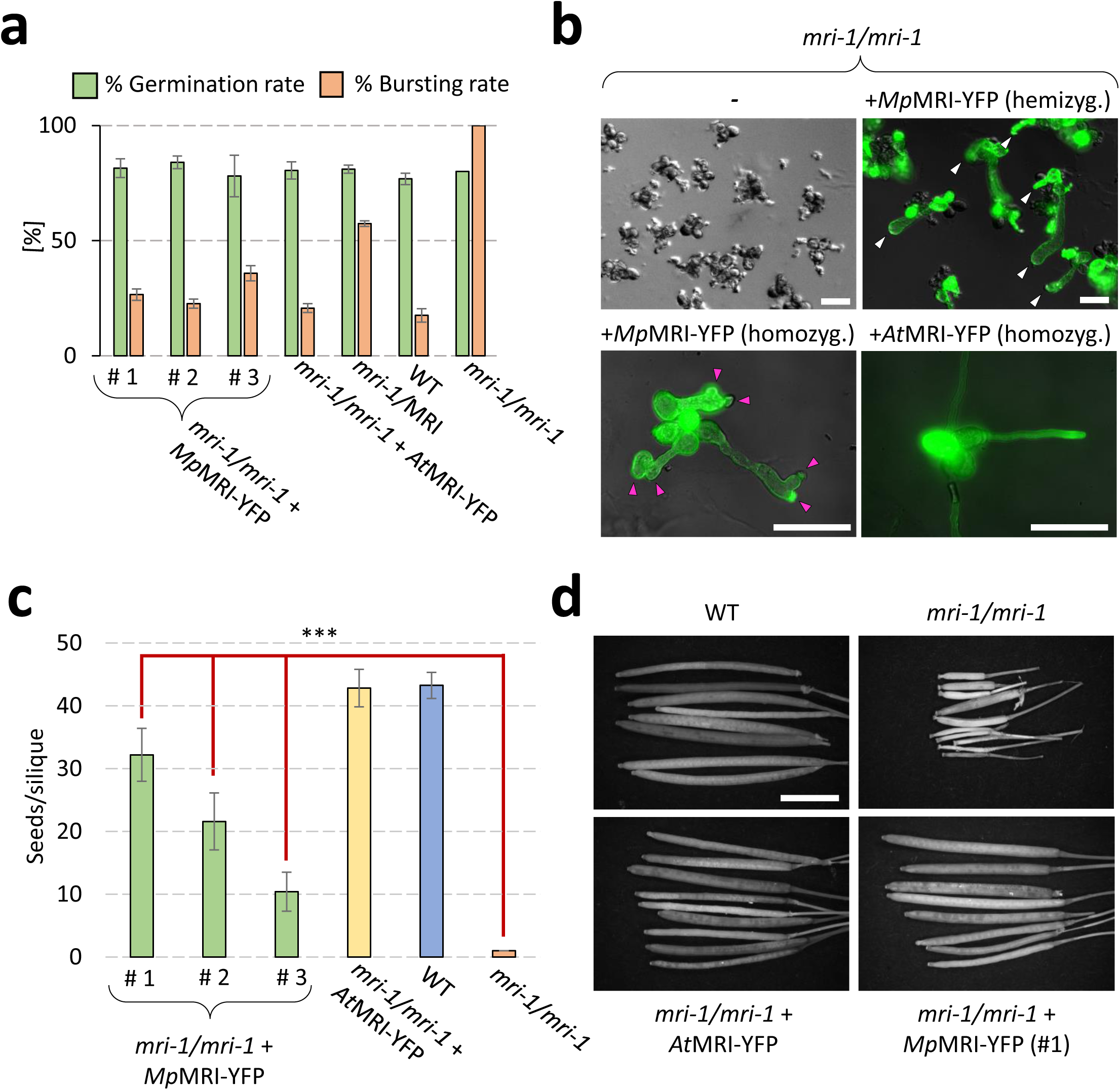
*Mp*MRI can rescue *mri-1*-pollen bursting and male sterility. **(a)** *In vitro* pollen germination assay of three independent T3 lines of *mri-1/mri-1* homozygously expressing *proAt*MRI:*Mp*MRI-YFP, in comparison to WT, *mri-1*/MRI, *mri-1/mri-1* and *mri-1/mri-1* complemented with *proAt*MRI:*At*MRI-YFP. Pollen bursting rates of all three *mri-1* lines expressing *Mp*MRI-YFP were not significantly different from *mri-1* expressing *At*MRI-YFP or WT (p>0.01 in an unpaired, two-tailed student’s T-test); a total of 600 pollen grains was analyzed in n = 3 independent experiments; error bars show the standard error of the mean. **(b)** Representative pictures of the data shown in section (a). White arrowheads indicate pollen tubes in a hemizygous *Mp*MRI-YFP expression line, all of which are transgenic. Pink arrowheads indicate an abnormal shape, *i.e.* pollen tube branching, swelling of tube apices and tip-focused lesions. Scale bars = 80 μm. **(c)** Seed set assay of individuals of three independent T3 lines of *mri-1/mri-1* expressing *proAt*MRI:*Mp*MRI-YFP, as compared to *mri-1/mri-1* complemented with *proAt*MRI:*At*MRI-YFP, WT and untransformed *mri-1/mri-1*. Datasets of all three transformant lines were significantly different from *mri-1/mri-1* (unpaired, two-tailed student’s T-test; ***: p<0.001; n = 15 siliques per individual). **(d)** Representative siliques of the plants analyzed in (c). Scale bar: 2 mm.

To test if expression of *Mp*MRI-YFP can rescue the loss of CWI in *mri-1* mutant root hairs, we analyzed root hair growth in two of the *A. thaliana mri-1* T3 lines homozygous for *Mp*MRI-YFP. Not only was root hair length partially restored to wild-type length in all individuals (n > 10; Fig. 4a,c), but the rate of root hair bursting was much lower than in *mri-1* mutants (Fig. 4b,c). *mri-1* root hairs expressing *Mp*MRI-YFP (Fig. 3b) developed cylindrical root hairs indistinguishable from wild type (Fig. 4c). *Mp*MRI-YFP localized to the plasma membrane of root hairs similarly as it and *At*MRI did in other tip-growing cells (Fig. 4c). These results demonstrate that *Mp*MRI can fully replace *At*MRI in the CWI maintenance pathway during root hair growth.

**Figure 4:**
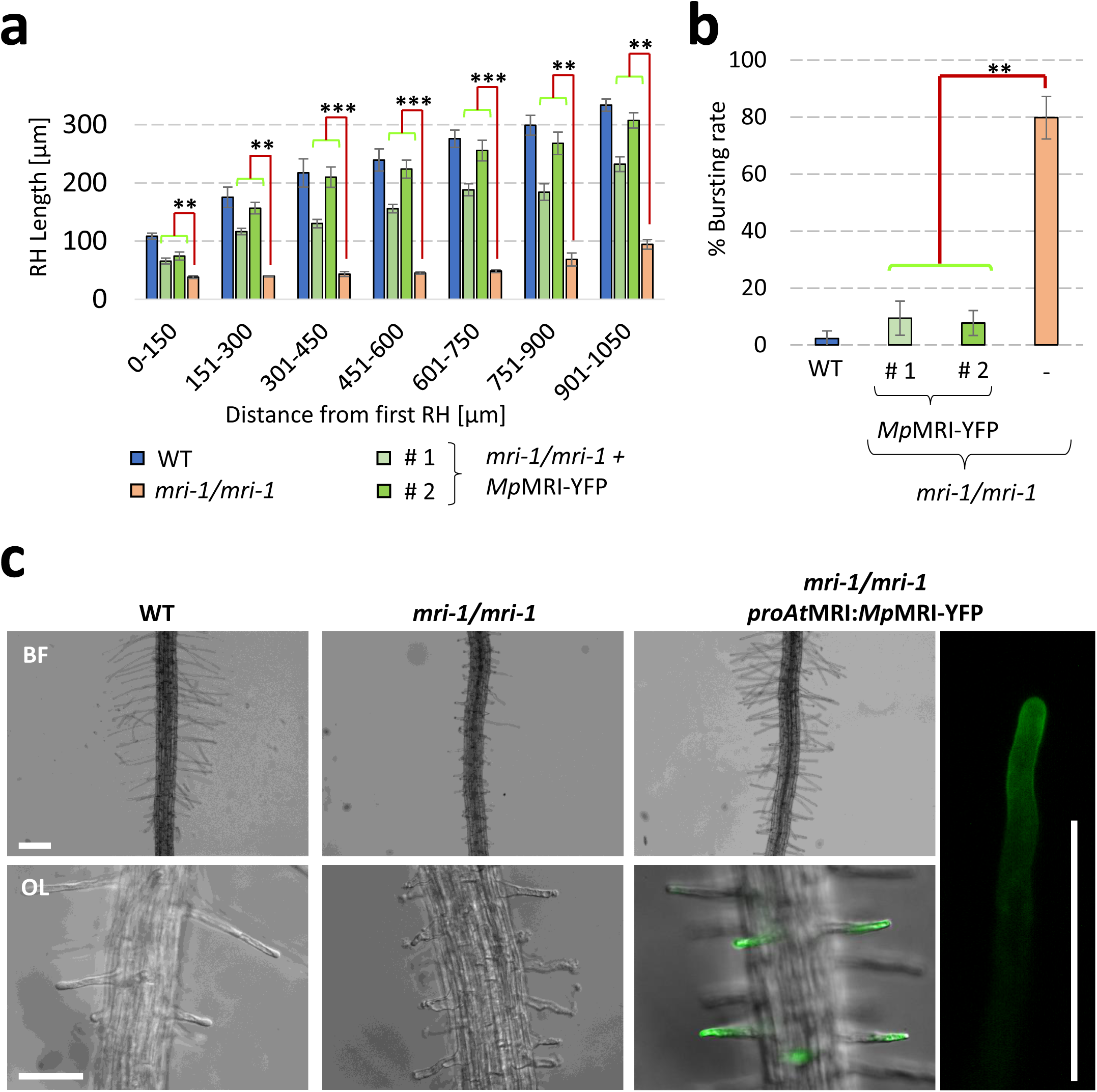
*Mp*MRI can maintain CWI in growing *mri-1* root hairs. **(a)** Root hair length profile of *mri-1* expressing *proAt*MRI:*Mp*MRI-YFP (n = 2 independent lines). Each primary root was digitally divided in sections of 150 μm and root hair length was determined for 5-10 root hairs per section. Shown are the mean values of n = 5 seedlings per genotype and section; error bars show the standard error from the mean. Significance tested with a two-tailed, unpaired student’s T-test; **: p<0.01, ***: p<0.001. **(b)** Mean root hair bursting rate of n ≥ 16 seedlings; error bars show the standard deviation from the mean value. Significance tested with a two-tailed, unpaired student’s T-test; **: p<0.01, ***: p<0.001. **(c)** Representative captures of the data presented in (a) and (b). BF = bright field; OL = Overlay of bright field image and plasma membrane *Mp*MRI-YFP derived fluorescence. Scale bar: 200 μm.

### Overexpression of PTI1-like proteins inhibits rhizoid, root hair and pollen tube growth

Overexpression of positive or knockdown of negative CWI regulators can lead to growth cessation in Arabidopsis pollen tubes and root hairs^16–18^. To further confirm that *Mp*MRI function is conserved between *A. thaliana* and *M. polymorpha* we tested whether rhizoid growth would be inhibited by overexpression of PTI1-like genes. Constitutively overexpressing *MpMRI* and *AtMRI* fused to a ternary Citrine tag (3xCitrine) in wild-type sporelings led to significant rhizoid growth inhibition (Fig. 5a,b and Supplementary Fig. 5a). The shorter rhizoids on lines overexpressing *MpMRI*- or *AtMRI*-*3xCitrine* displayed the expected plasma membrane localization for both *Mp*MRI and *At*MRI (Supplementary Fig. 3). Taken together, these results indicate that the repression of growth by PTI1-like kinases has been conserved since *A. thaliana* and *M. polymorpha* last shared a common ancestor.

**Figure 5:**
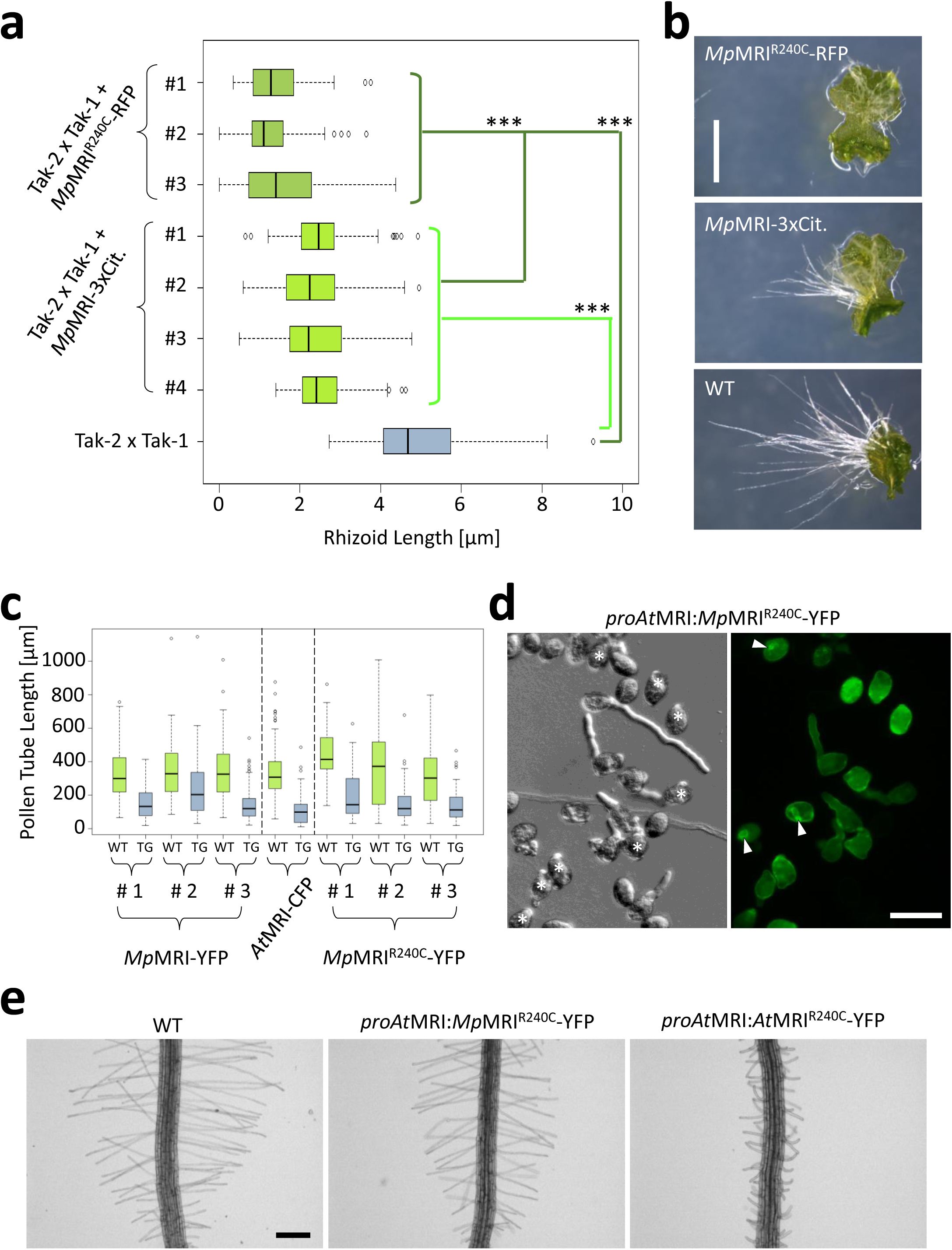
Expression of PTI1-like receptor-like cytoplasmic kinases triggers growth inhibition in Marchantia and Arabidopsis tip-growing cells. **(a)** Rhizoid length of 7 d old Marchantia Tak-2 x Tak-1 wild-type thalli expressing *Mp*MRI, or *Mp*MRI^240C^, respectively, compared to a non-transformed Tak-2 x Tak-1 reference line (n > 10 thalli/independent line). Significance of all expression lines against non-transformed Tak-2 x Tak-1 and all *Mp*MRI^240C^ lines against *Mp*MRI lines, was tested with a two-tailed, unpaired student’s T-test (***: p<0.001). See also Supplementary Fig. 5a for *At*MRI expression lines. **(b)** Representative images of the data shown in (a). Scale bar: 2 mm. **(c)** *In vitro* pollen tube length measurements of Arabidopsis wild-type lines hemizygously expressing *proAt*MRI:*Mp*MRI-YFP (n = 3 lines), *pro*Lat52:*At*MRI-CFP or *proAt*MRI:*Mp*MRI^R240C^-YFP (n = 3 lines), respectively. The 10 longest wild-type (WT) and transgenic (TG) pollen tubes per image were measured, n ≥90 pollen tubes per line. Significance was tested with an unpaired, two-tailed student’s T-test (***: p<0.001; see also Supplementary Fig. 5b). **(d)** Representative captures of the data presented in section (c). Note how most of the fluorescent pollen only form bulges (asterisks) and occasionally some plasma membrane invaginations (arrowheads). Scale bar = 50 μm. **(e)** Representative images of roots of wild type and *Mp*MRI^R240C^-YFP and *At*MRI^R240C^-YFP expressing seedlings. For quantification of root hair lengths see Supplementary Fig. 6; scale bar: 200 µm.

### The conserved R240 of PTI1-like proteins exerts an auto-inhibitory effect

A suppressor screen for rescue of male fertility in the double-knockout mutant *Atanx1 anx2* helped identify a hyperactive variant of *At*MRI, with a R240C substitution in the activation loop of the kinase domain^18^. Structural similarity analyses of land plant PTI1-like revealed that *Mp*MRI is part of a monophyletic group that includes *At*MRI (termed the ‘MRI-like group’ from here on; Supplementary Fig. 4a). Both the invariant lysine residue of the ATP-binding pocket required for kinase activity (K100 in *At*MRI) and the STR-motif, which includes arginine R240, are conserved among members of the MRI-like group (Supplementary Fig. 4a). Furthermore, the conserved threonine (T239 in *At*MRI) of the conserved STR motif of some PTI1-like kinases can be phosphorylated by other kinases such as the Arabidopsis OXIDATIVE SIGNAL-INDUCIBLE1 (OXI1) and tomato Pto kinases^25, 26^.

To better understand the mechanistic role of conserved K100, T239 and R240 residues, we designed several structural variants of *At*MRI (Supplementary Table 3). First, we designed two kinase-dead variants with a K100N and K100E amino acid substitution, respectively ^21, 27, 28^ and introduced them in *mri-1*/MRI under the pollen-specific promoter *Lat52*^29^. Expression of both structural *At*MRI^K100E^ and *At*MRI^K100N^ variants led to the emergence of homozygous *mri-1/mri-1* lines in the self-progeny of transformed *mri-1*/*MRI* (Supplementary Table 3, orange rows) similarly as for expression of wild-type *At*MRI (Supplementary Table 3, green row) or *At*MRI^R240C^ (Supplementary Table 3, blue row). Hence, kinase activity does not appear to be required for *At*MRI function during pollen tube growth. Second, we tested for the necessity of phosphorylation at residue T239 by engineering a non-phosphorylatable T239A and a phosphomimetic T239E variant, respectively, and introducing them in *mri-1*/*MRI*. Again, expression of both *At*MRI^T239A^ and *At*MRI^T239E^ variants led to the emergence of homozygous *mri-1/mri-1* individuals (Supplementary Table 3, yellow rows) indicating that phosphorylation at T239 is not required for *At*MRI function during pollen tube growth. These results are in good agreement with a previous study on rice *Os*PTI1a, where equivalent mutant variants *Os*PTI1a^K96N^ and *Os*PTI1a^T233A^ were able to rescue *Ospti1a* spontaneous lesions and rice blast-enhanced resistance phenotypes^30^. Moreover, they also suggest that overactivity of *At*MRI^R240C^ arises from a substitution of R240 itself rather than from mimicking phosphorylation at T239^31^. We then produced some variants for R240 itself, namely R240M, R240A and R240S. To assess a potential protein overactivity that may arise from the substitutions, we expressed all our variants in the pollen tube bursting male-sterile *AtrbohH rbohJ* mutant, whose sterility is rescued by expression of the overactive *At*MRI^R240C^ variant, but not the native *At*MRI protein^18^. Similarly, as for wild-type *AtMRI*, expression of the phosphomimetic *AtMRI^T239E^*variant did not restore *AtrbohH rbohJ* sterility, with all independent T1 lines producing only a few seeds (Supplementary Fig. 4b). However, expression of any R240 variant led to a clear rescue of *AtrbohH rbohJ* fertility with seed set increase (Supplementary Fig. 4b). These results confirm that R240C overactivity is not due to phospho-mimicking at residue T239 and indicate that any substitution of residue R240 leads to an overactive version of *At*MRI. Thus, residue R240 of *At*MRI appear to exert an enigmatic auto-inhibitory effect.

### *Mp*MRI^R240C^ inhibits growth in Marchantia and Arabidopsis tip-growing cells

If *Mp*MRI and *At*MRI are functionally conserved, we would expect that the hyperactive R240C mutation in each would have similar phenotypic consequences. To test this, we constitutively expressed variant *Mp*MRI^R240C^-RFP in Marchantia. Rhizoid length was significantly reduced in all independent transformed lines compared to controls (Fig. 5a, b). Also, the growth inhibitory effect derived from *MpMRI^R240C^*expression was stronger than for *MpMRI* expression as observed when *AtMRI^R240C^* and *AtMRI* are overexpressed in *A. thaliana*^18^, suggesting that the R240C substitution in *Mp*MRI^R240C^ induces a similar effect as for *At*MRI^R240C^.

Next, to test whether *Mp*MRI^R240C^ growth inhibition effect is restricted to rhizoid, we introduced *Mp*MRI^R240C^-YFP in Arabidopsis wild-type pollen tubes and root hairs under the control of *proAt*MRI. To directly assess the effect of *Mp*MRI^R240C^ on pollen tube germination and growth, we used several independent lines hemizygously expressing the transgene, i.e. in plants from which half of the pollen is wild type and the other half expresses *MpMRI^R240C^-YFP* (Supplementary Table 4), thus allowing direct comparison of transgenic and wild-type pollen tubes within the same line and assays. Pollen tubes were shorter for pollen expressing *Mp*MRI^R240C^ than for wild-type pollen (Fig. 5c and d; Supplementary Fig. 5). We then asked whether these growth inhibitory effects were strong enough to affect pollen germination itself. While *Mp*MRI-YFP did not significantly affect the percentage of transgenic to non-transgenic pollen tubes (expected to be 50% if the transgene does not influence germination), expression of *Mp*MRI^R240C^-YFP decreased the ratio of pollen tubes carrying the transgene to 37 - 43 % (Supplementary Table 4). Interestingly, plasma membrane invaginations in fluorescent non-germinating pollen grains were observed, a phenotype described upon *At*MRI^R240C^ expression, as well (Fig. 5d)^18^. A more severe effect was observed upon expression of *At*MRI-CFP (16 %), while *At*MRI^R240C^-CFP induced a complete inhibition of pollen germination (complete absence of fluorescent pollen tubes), just as reported before (Supplementary Table 4)^18^. These findings suggest that expression of *Mp*MRI^R240C^ disrupts pollen tube growth more than *Mp*MRI as observed upon expression of *At*MRI^R240C^ and *At*MRI.

We assessed whether expression of *Mp*MRI^R240C^ has a similar growth inhibitory effect on root hairs. Root hairs were shorter on all lines homozygous for *proAt*MRI:*Mp*MRI^R240C^-YFP (originating from the same three independent T1 lines tested for pollen tube growth) than on wild type (Fig. 5e and Supplementary Fig. 6). Again, the growth inhibitory effect observed in root hairs was less pronounced than upon expression of Arabidopsis *At*MRI^R240C^-YFP (two independent lines, Fig. 5e and Supplementary Fig. 6). Altogether, the similar growth inhibitory effects observed on tip-growing Marchantia and Arabidopsis cells upon expression of either *Mp*MRI^R240C^ or *At*MRI^R240C^ suggest functional conservation of the auto-inhibitory residue R240 and overactivation of the CWI pathways upon expression of the R240C variants.

### *Mp*MRI and *Mp*FER act in a common, evolutionarily conserved signaling pathway

We previously reported that expression of *AtMRI^R240C^*was sufficient to partially rescue loss of CWI of *Atanx1 anx2* pollen tubes as well as *Atfer* root hairs, thereby positioning MRI downstream of the MLRs *At*ANX1 and *At*FER^18^. Interestingly, disruption of the unique Marchantia MLR homologue *Mp*FER^24^ was also reported to trigger loss of CWI in rhizoids^23^ suggesting that the MLR/PTI1-like signaling module is conserved during rhizoid growth. To further demonstrate the conservation, we expressed a *Mp*MRI^R240C^-RFP protein fusion in the *Mpfer-1* mutant^23^ (Supplementary Fig. 1). In contrast to *Mpfer-1* in which almost every rhizoid bursts at an early stage of growth, rhizoids on 4 independent *Mpfer-1* lines transformed with *Mp*MRI^R240C^-RFP were longer and more frequently intact (Fig. 6 and Supplementary Fig. 7). These data indicate that *Mp*MRI functions downstream of *Mp*FER in the CWI pathway that controls rhizoid growth. They also confirm that the MLR/PTI1-like signaling module controlling CWI in tip-growing cells has been conserved during land plant evolution from the rhizoids of liverworts to the root hairs and pollen tubes of seed plants.

**Figure 6:**
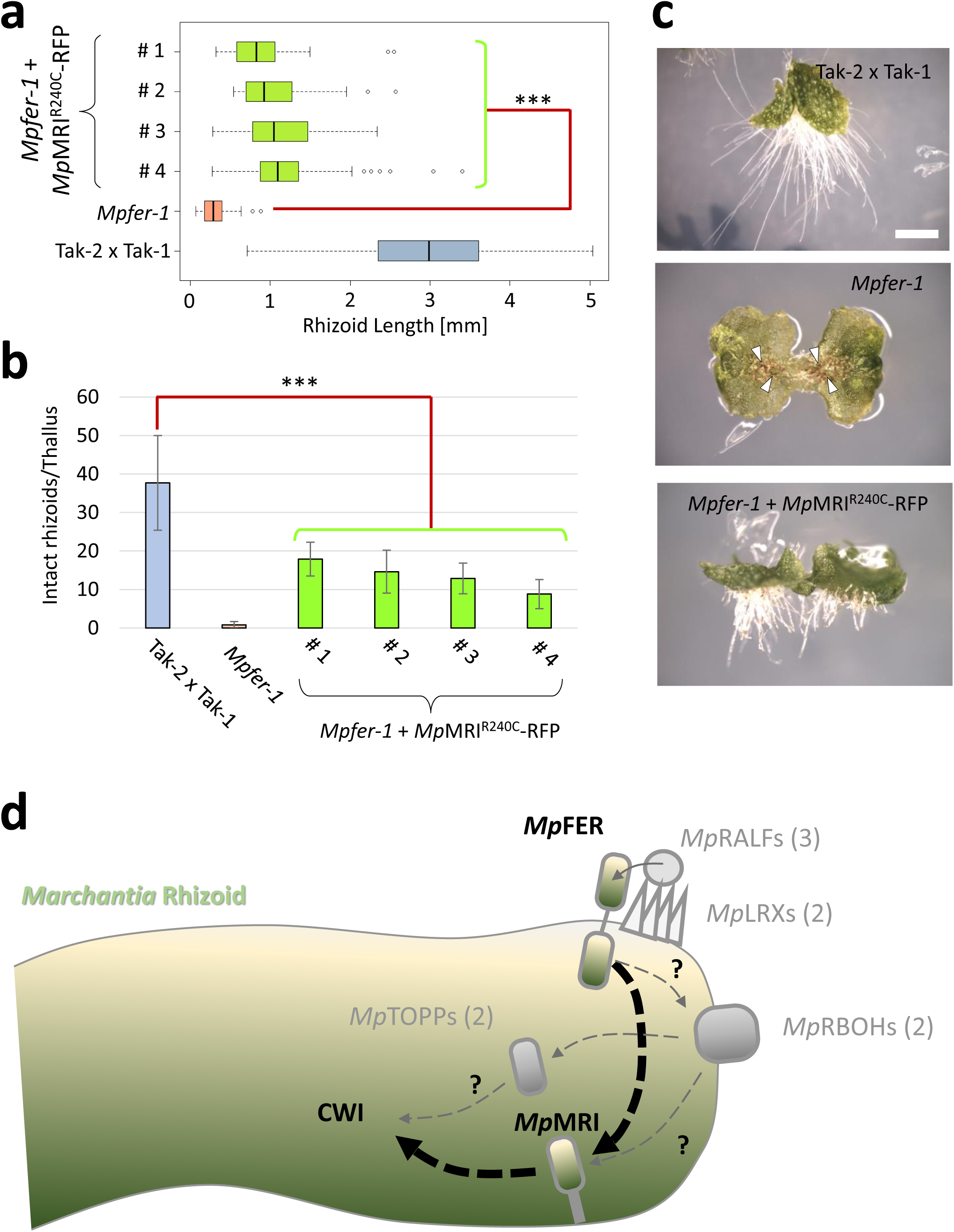
*Mp*MRI functions downstream of *Mp*FER to maintain CWI in growing rhizoids. **(a)** Rhizoid length of 7 d old *Mpfer-1* thalli transformed with *proMp*EF1α:*Mp*MRI^R240C^-RFP (n = 4 independent lines) compared to Tak-2 and non-transformed *Mpfer-1* (n > 10 thalli/ independent line). Significance of all expression lines against non-transformed *Mpfer-1* was tested with a two-tailed, unpaired student’s T-test (***: p<0.001). **(b)** Number of intact rhizoids of 7 d old *Mpfer-1* thalli transformed with *proMp*EF1α:*Mp*MRI^R240C^-RFP (n = 4 lines) compared to Tak-2 and non-transformed *Mpfer-1* (n ≥ 10 individuals/ genotype). Significance of all 4 transformed lines against non-transformed *Mpfer-1* was tested with a two-tailed, unpaired student’s T-test (***: p < 0.001). **(c)** Representative images of the lines analyzed in sections (a) and (b). Arrowheads depict the frequent loss of CWI at the tip of *Mpfer-1* rhizoids, which is decreased in all *Mpfer-1* lines transformed with *Mp*MRI^R240C^-RFP. Scale bar: 1 mm. See also Supplementary Fig. 7 for older thalli. **(d)** Working model of the genetic rhizoid CWI control pathway: The Malectin-like receptor *Mp*FER controls CWI maintenance in the growing Marchantia rhizoid through *Mp*MRI. Whether activation and signal transduction are mediated via similar signaling constituents (shown in grey) as in Arabidopsis remains to be tested. Numbers in brackets indicate the number of respective Marchantia homologs.

## Discussion

During land plant evolution, tip-growth was recruited to participate in several key adaptations to terrestrial habitats that facilitated colonization and radiation of land plants. Among these adaptations are substrate anchorage, water/nutrient uptake and microorganism interactions via rooting cells (rhizoids and root hairs), and water-independent sperm delivery to the egg cell via pollen tubes. While tip-growth in plants performs extremely well for increasing surfaces, exploring environment and transport, it relies on a delicate balance of turgor-driven, anisotropic cell expansion and appropriate secretion of surface membrane and cell wall material that defines cell shape and function. We show that this balance is mediated, at least in part, by the FER/MRI signaling module in tip-growing cells. Failure of this mechanism in plants with defective FERONIA or MARIS function results in loss of cellular integrity because of mechanical failure in the cell wall.

While rhizoids and root hairs represent morphologically similar cell types serving similar biological functions, they develop during different life-phases. Rhizoids are found on the gametophyte of early diverging land plant taxa, while root hairs develop on the roots of vascular plants^4, 32^. The development of rhizoids or root hairs in all major lineages of land plants suggests that tip-growing cells formed the rooting structures of the first land plants^4^. Unlike pollen tubes that emerge from pollen grains, rhizoids and root hairs originate from the swelling of some epidermal cells. The development of such is controlled by an ancient conserved genetic mechanism composed of the class I ROOT HAIR DEFECTIVE SIX-LIKE (RSL) basic helix-loop-helix transcription factors^33, 34^. The control of root hair and rhizoid development by the same transcription factors has been interpreted as an example of deep homology^35^ since, at the level of land plants, rhizoids and root hairs are analogous structures with similar functions whose development is controlled by a homologous genetic mechanism.

Later during evolution, some vascular plants developed an elaborate system to guarantee an efficient water-independent sperm cell delivery to the female egg cell via pollen tubes, a complex process referred to as siphonogamy^5, 36^. Siphonogamy may have evolved twice independently within the gymnosperm and angiosperm lineages^5, 37^. Pollen tubes of seed plants are thought to have evolved from a slow and long-lived haustorial pollen tube involved in nutritional uptake to a fast-growing, short-lived pollen tube whose unique function is the delivery of non-motile sperm in angiosperms^5, 38^. It is currently unknown whether CWI maintenance mechanisms were already active in the slow haustorial pollen tubes or if they were recruited along the transition to fast growth to avoid growth accidents and infertility. So far, however, there have been no reports of homologous mechanisms conserved between pollen tubes and rhizoids.

In this study, we demonstrate the evolutionarily conservation of the genetic pathway comprised of MLRs and PTI1-like receptor-like kinases that regulates CWI maintenance during tip-growth. We showed that not only disruption but also overexpression of the PTI1-like encoding genes *MpMRI* and *AtMRI* led to similar CWI maintenance defects in rhizoids of the early diverging land plant Marchantia as in the root hairs and pollen tubes of the seed plant Arabidopsis. Moreover, trans-species complementation assays showed that *AtMRI* expression partially rescues the bursting rhizoid phenotype of *Mpmri-1* while *MpMRI* expression rescues partially pollen tube bursting of *Atmri-1* and fully its bursting root hair phenotype. These results demonstrate that *Mp*MRI and *At*MRI can largely functionally replace each other regardless of the tip-growing cell type or the phylogenetic position of the plant on which they develop. This indicates that the fundamental function of PTI1-like proteins has been conserved since liverworts and seed plants diverged from a common ancestor. Nonetheless, in the Arabidopsis *mri-1* mutant, *Mp*MRI-expressing root hairs grew better than *Mp*MRI-expressing pollen tubes. Consequently, the CWI control in sporophytic root hairs is likely to be more similar to that in gametophytic rhizoids than in pollen tubes, the male gametophytes of flowering plants. Thus, our study suggests that a genetic program maintaining CWI in rhizoids was established early during land plant evolution and then adopted by the sporophyte for root hair growth control and much later by the male gametophyte of flowering plants.

One can also wonder whether CWI maintenance during tip-growth represents the original ancestral function of PTI1-like proteins. The fact that several Arabidopsis PTI1-like proteins, close homologues of *At*MRI, cannot even partially substitute for *At*MRI in pollen tube CWI maintenance^39^ but *Mp*MRI can, suggests subfunctionalisation of the PTI1-like gene family in the lineage that led to Arabidopsis. Studies in tomato, cucumber, and rice all demonstrate a role for PTI1-like in plant responses to pathogens as either positive (in dicots such as Tomato^21, 40^ and Cucumber^41^) or negative (in monocot, Rice^28, 42^) regulators. Moreover, the Arabidopsis PTI1-like *At*CARK1 positively regulates abscisic acid (ABA) signaling by phosphorylating ABA receptors^22^. Since some ABA responses and signaling components are conserved in Marchantia^24, 43–45^, and pathosystems are being established in Marchantia^46^ it would be informative to test if *Mp*MRI regulates ABA signaling and/or immunity in this species. Our previous study showed that the *At*MRI^R240C^ structural variant functions as an overactive protein version, whose expression is not only sufficient for tip-growth inhibition in the wild-type background but also rescues loss of CWI in mutants of its upstream MLR regulators *At*ANX1/2 for pollen tubes and *At*FER for root hairs^18^. Here, we showed that expression of the Marchantia variant *Mp*MRI^R240C^ triggered similar growth inhibitory effects in both, Marchantia and Arabidopsis tip-growing cells suggesting a conserved structural mechanism. Furthermore, expressing the *Mp*MRI^R240C^ variant in the *Mpfer* mutant partially rescued *Mpfer* bursting rhizoids, similarly as did *At*MRI^R240C^ expression for *Atfer* bursting root hairs and *Atanx1 anx2* bursting pollen tubes. Therefore, the distantly related land plant taxa Arabidopsis and Marchantia share a common MLR/PTI1-like receptor-like kinase signaling module to control CWI in tip-growing cells. Since these two species last shared a common ancestor approximately 450-470 million years ago and given the most likely topologies of land plant phylogeny, we conclude that such a signaling module is an ancient common feature shared by most lineages of extant land plants. Whether such signaling module includes further common regulatory features known from the Arabidopsis CWI maintenance pathways (Fig. 6d), such as extracellular receptor binding of RALF peptides, regulation of ROS-producing NADPH oxidase etc. remains to be investigated.

## Materials and Methods

### Plant materials and growth conditions

All mutant lines, transgenic lines, oligonucleotide sequences and plasmids used in this study are listed in Supplementary Tables 5-8, respectively. The presence and position of the T-DNA in the respective genetic locus of the insertional mutants *Mpmri-1* (ST13.3) and *Mpfer-1* (ST14.3)^23^ was verified via PCR-genotyping and sequencing (Supplementary Fig. 1). Binary vectors were first transformed into *Agrobacterium tumefaciens* strain GV3101 via electroporation. A pure culture was then used to transform Arabidopsis^47^ or Marchantia sporelings^48^ and thalli^49^.

Arabidopsis seeds (Col-0) were sown on solid half-strength Murashige and Skoog (MS) basal medium (Duchefa Biochemie), stratified (2 d, 4 °C, darkness) and allowed to germinate in a growth chamber set to long day conditions (7-10 d, 21 °C, 60 % humidity, 16 h white light at 80 µmol m^-1^ s^-1^/8 h darkness cycle). Seedlings were then transferred and grown on soil in a greenhouse under long day conditions. Vegetative Marchantia stages were grown on solid Johnson’s growth medium^50^ (0.8 % agar to facilitate growth of rhizoid defective mutants) under long day conditions. To cross male and female Marchantia lineages, 4 weeks old thalli were transferred onto soil and grown for 3 weeks under long day conditions (white light: 40 µmol m^-1^ s^-1^; far-red light between 700-880 nm: 15 µmol m^-1^ s^-1^) to trigger gametangia formation. Crossings were then carried out manually via spermatozoid transfer from mature male to female gametangia. Emerging sporangia were harvested after 2-3 weeks and sterilized with 1 % sodium dichlorisocyanurate to allow for axenic, vegetative cultivation.

### Molecular cloning

To amplify the *Mp*MRI and *Mp*FER coding sequences (CDS), RNA was isolated from Tak-1 and Tak-2 whole-plant material using the Direct-Zol RNA Mini-Preparation Kit (Zymo Research). cDNA synthesis was carried out using the RevertAid H Minus First Strand cDNA Synthesis Kit (Thermo Fisher Scientific) and used as template for the amplification of the respective CDS without stop codon using gateway-compatible primers. The different CDS were then cloned into pDONR207 and transformed into *Escherichia coli DH5α* cells to generate Gateway-entry clones. The CDS were then remobilized in the respective Gateway destination vectors of choice to generate fluorescent protein fusions (Supplementary Table 7). Amplified sequences and preservation of the reading frames were verified via sequencing of the full-length CDS (entry clones) and the promoter-CDS / CDS-fluorescent protein junctions (final binary constructs).

The new variants *Mp*MRI^R240C^ and *At*MRI^K100N^, *At*MRI^T239A^, *At*MRI^T239E^, *At*MRI^R240M^, *At*MRI^R240A^ and *At*MRI^R240S^, were generated via site-directed mutagenesis (SDM). SDM primers were designed using Primer X (http://www.bioinformatics.org/primerx/) to introduce via PCR the different adequate nucleotide substitutions with *Mp*MRI and *At*MRI in pDONR207 as template. The PCR-mix were digested with the methylation-sensitive restriction enzyme *Dpn*I to degrade non-mutagenized vector copies. The mutagenized vectors were transformed into *E. coli DH5α* cells and the respective nucleotide substitution were verified via sequencing before remobilization of the Gateway cassette in the final binary plasmids.

### Selection of transformant lines

Putatively transformed Arabidopsis T1 seeds were preselected on solid half-strength MS with Basta (glufosinate ammonium, 10 µg/ mL). Resistant T1 seedlings were transferred and grown on soil. Upon flower emergence, pollen was checked for expression of the respective fluorescent fusion protein. Transgenic T1 lines with clear, homogenous fluorescence in about 50 % of the pollen were propagated. Their seeds were harvested and selected again in the T2 generation. Upon flower emergence, plants were screened for 100 % fluorescent pollen grains, indicating homozygous lines. Plants expressing fluorescent protein fusions under control of *proAt*MRI were additionally screened for root hair-specific fluorescence at the seedling stage.

Putatively transformed Marchantia sporelings/thalli were preselected on solid Johnson’s growth medium with Chlorsulfuron (100 µg/ mL) or Gentamicin (0.5 µM) and supplemented with Cefotaxime (100 µg/ mL) to avoid contamination with residual Agrobacteria. Resistant plants were then genotyped for presence of the fluorescent protein fusion and additionally checked for Citrine/RFP fluorescence in thalli, gemmae and/or rhizoids. Propagation of the different transgenic lines was done by transplantation and cultivation of gemmae.

### Rhizoid growth assays

Gemmae were grown vertically for 7 d under axenic standard conditions (as described above). To determine the mean rhizoid length, the 10 longest rhizoids per gemmaling were measured for n ≥ 10 individuals. To approximately estimate the percentage of bursting rhizoids, the mean number of visibly intact rhizoids per thallus was determined per individual. For time course experiments, data was collected after 7, 14, 21 and 28 d. Images were taken with a Leica MZ 16 F fluorescence binocular (whole-plant), a Leica DM5000 fluorescence microscope (close-up images). Alternatively, fresh gemmaes were grown for 24hr in liquid Johnson’s growth medium sandwiched between a slide and a cover slip separated with a thin layer of parafilm. Young growing rhizoids were then imaged with either a Leica DM5000 fluorescence microscope or a Leica SP8 confocal laser scanning microscope.

### *In vitro* Pollen germination assays and length measurements

Young, freshly opened Arabidopsis flowers were harvested in the morning and pollen grains were allowed to hydrate for 30 min under long day conditions in a humid environment. Stamina were then brushed on a microscopic slide coated with 500 µL of solid pollen germination medium ^51^. Pollen germination assays and imaging were conducted as described previously^17^. Bright field microscopy was carried out with a Leica DM5000 microscope and n = 150 – 300 pollen grains per genotype and experiment were imaged and later classified as non-germinated, germinated and intact or germinated and burst.

In order to determine the mean pollen tube length, pollen germination was induced as described above. The length of n = 200 pollen tubes was measured following the grown pollen tube from the origin of germination to the pollen tube tip in three independent experiments.

### Seed set assays

Plants were allowed to self-fertilize and n > 10 mature siliques were harvested per individual plant. Siliques were fixed and bleached in 1.5 mL of an acetic acid / ethanol solution (v/v 1:3) for 1 d. Siliques were then opened and their seeds were quantified to determine the mean number of seeds per silique. Images were obtained with a Leica MZ 16 F fluorescence binocular.

### Root hair assays

Seeds were allowed to germinate vertically on solid half-strength MS medium supplemented with 2 % sucrose. 4 – 5 d old seedlings were embedded in a microscope slide with liquid 1/10 MS medium. The borders of the glass slide were wrapped with a thin parafilm layer to act as spacer to prevent squashing the young fragile Arabidopsis roots with the coverslip. The most distal part of the root was imaged (Leica DM5000 fluorescence microscope) and digitally divided into sections of 150 µm length starting from the first root hair from the tip. The length of 5 – 10 root hairs per section was determined. The mean of each section was then compared amongst specimens of each genotype. The root hair bursting rate was determined as the percentage of bursting root hairs out of the total root hair number.

### Phylogenetic analyses

The AtMRI (AT2G41970) amino acid sequence was used as query to perform a pBLAST homology search (BLOSUM62 matrix) against the protein database on representative species of major land plant taxa. pBLAST was performed on https://phytozome.jgi.doe.gov/ using standard settings for all species. The following species and genome versions were used: AmTr - *Amborella trichopoda* (v1.0), AT - *Arabidopsis thaliana* (TAIR10), Mapoly - *Marchantia polymorpha* (v3.1), Pp - *Physcomitrella patens* (v3.3), SeMo - *Selaginella moellendorffii* (v1.2). Only sequences either showing a percent identity >60% and/or annotation as PTO-interacting protein 1-like proteins (KEGG ORTHOLOGY code K13436) were considered. Multiple sequence alignments of all PTI1-like amino acid sequences were built using ClustalX2 ^52^ and illustrated using GeneDoc 2.7.0 and phylogenetic trees were calculated with MEGA7^53^ using the Maximum Likelihood method based on the JTT matrix-based model^54^ and 1000 repetitions. The tree with the highest log likelihood (-4460.1659) is shown in Supplementary Fig. 4. The percentage of trees in which the associated taxa clustered together is shown next to the branches. The frequency of amino acid usage per residue was illustrated using WebLogo 3 (http://weblogo.threeplusone.com/create.cgi).

### Data processing

Image analysis was performed using FIJI/ImageJ 1.50e^55^. Bar graphs were designed using Excel/Microsoft Office 365. Boxplots were designed using R studio 0.99.892 (RStudio Team, 2015).

## Acknowledgements

We thank Ueli Grossniklaus and Moritz Rövekamp (University of Zürich, Switzerland) for sharing unpublished data and enriching discussion. We also thank all members of Martin Hülskamp’s, Ute Höcker’s (University of Cologne, Germany) and Liam Dolan’s (University of Oxford, United Kingdom) groups for sharing their facilities. We thank Gabi Meineke (University of Cologne, Germany) for administrative support, Hiroyasu Motose (University of Okayama, Japan) and Benjamin Jaegle (Gregor Mendel Institute, Austria) for valuable intel about live-imaging of Marchantia rhizoids and phylogenetic trees, respectively. This research was partly funded by a short-term stipend of the Deutscher Akademischer Austauschdienst (DAAD) to J.W.; Biotechnology Biology Research Council Doctoral Train Partnerships fellowship in Interdisciplinary Bioscience to S.S.; European Research Council Advanced Grants EVO-500 (250284) and DENOVO-P grant (787613) to L.D; the University of Cologne, the Deutsche Forschungsgemeinschaft Grant BO 4470/1-1, and grant from the University of Cologne Centre of Excellence in Plant Sciences to A.B.-D..

## Author contributions

J.W., L.D. and A.B.-D. conceived the experiments. J.W., S.S., C.M.F., R.L., and A.B.-D. performed the experiments shown. J.W. and A.B.-D. analyzed the data. J.W., L.D. and A.B.-D. wrote the manuscript with contributions from the other authors.

## Competing interests

The authors declare no competing interests.

---

**Supplementary Video 1: Time course of a growing *Mpmri-1* rhizoid expressing *MpMRI-RFP*.**

Δt =15 sec, 30 frames. Left side is an overlay of RFP-derived fluorescence in magenta and plastids autofluorescence in green. Right side is bright-field. Scale bar: 10μm.

**Supplementary Video 2: Time course of a growing *Mpmri-1* rhizoid expressing *AtMRI-RFP*.**

Δt =15 sec, 30 frames. Left side is an overlay of RFP-derived fluorescence in magenta and plastids autofluorescence in green. Right side is bright-field. Scale bar: 10μm.

**Supplementary Figure 1:**
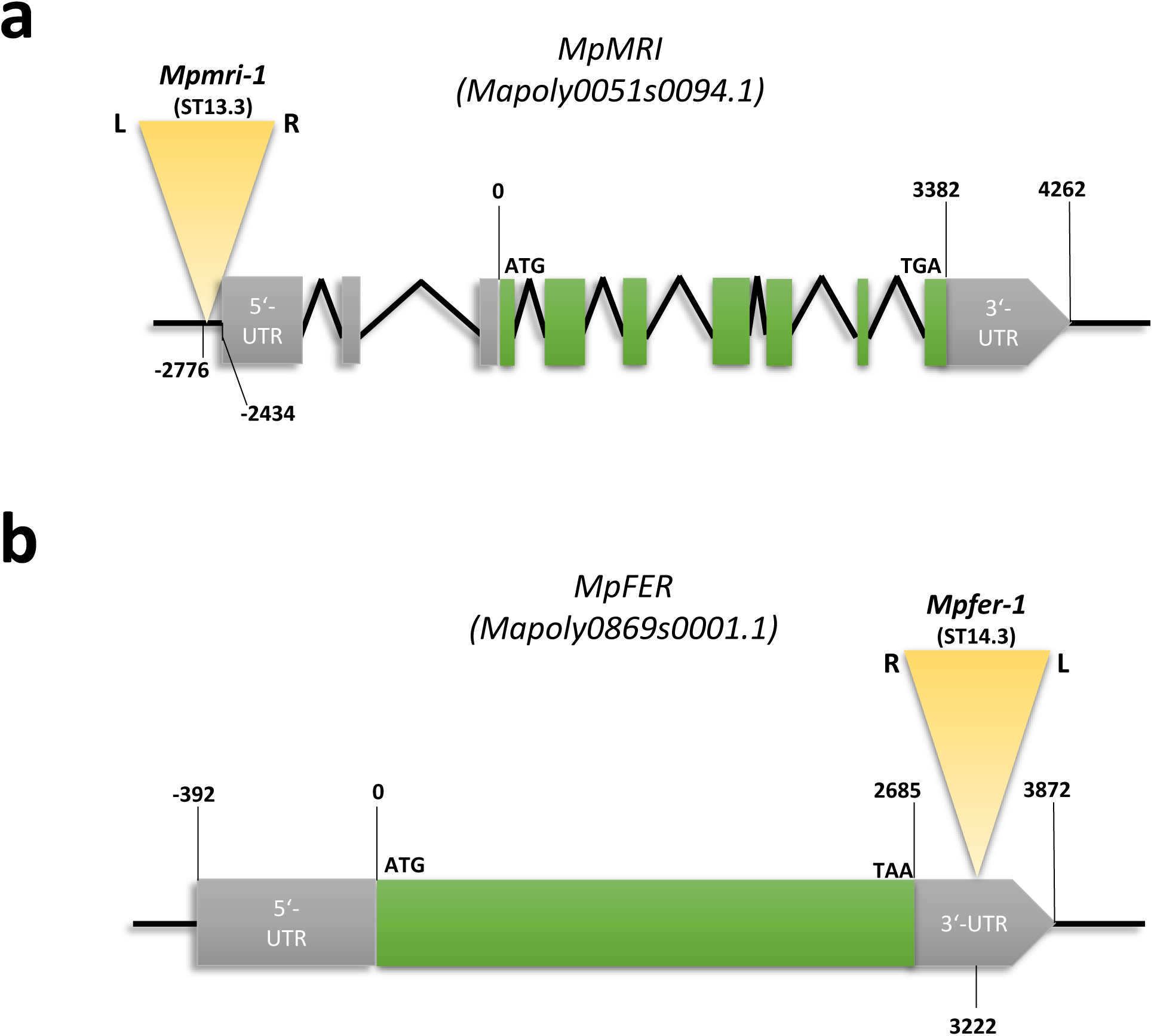
*MpMRI* and *MpFER* gene models and T-DNA insertion sites. **(a)** *MpMRI* (Mapoly0051s0094.1) and **(b)** *MpFER* (Mapoly0869s0001.1) schematic gene models; *Marchantia polymorpha* genome version 3. Introns, exons, untranslated regions (UTRs) and T-DNAs are depicted as tilted black lines, green boxes, grey boxes and yellow triangles respectively. All nucleotide positions refer to the start of translation.

**Supplementary Figure 2:**
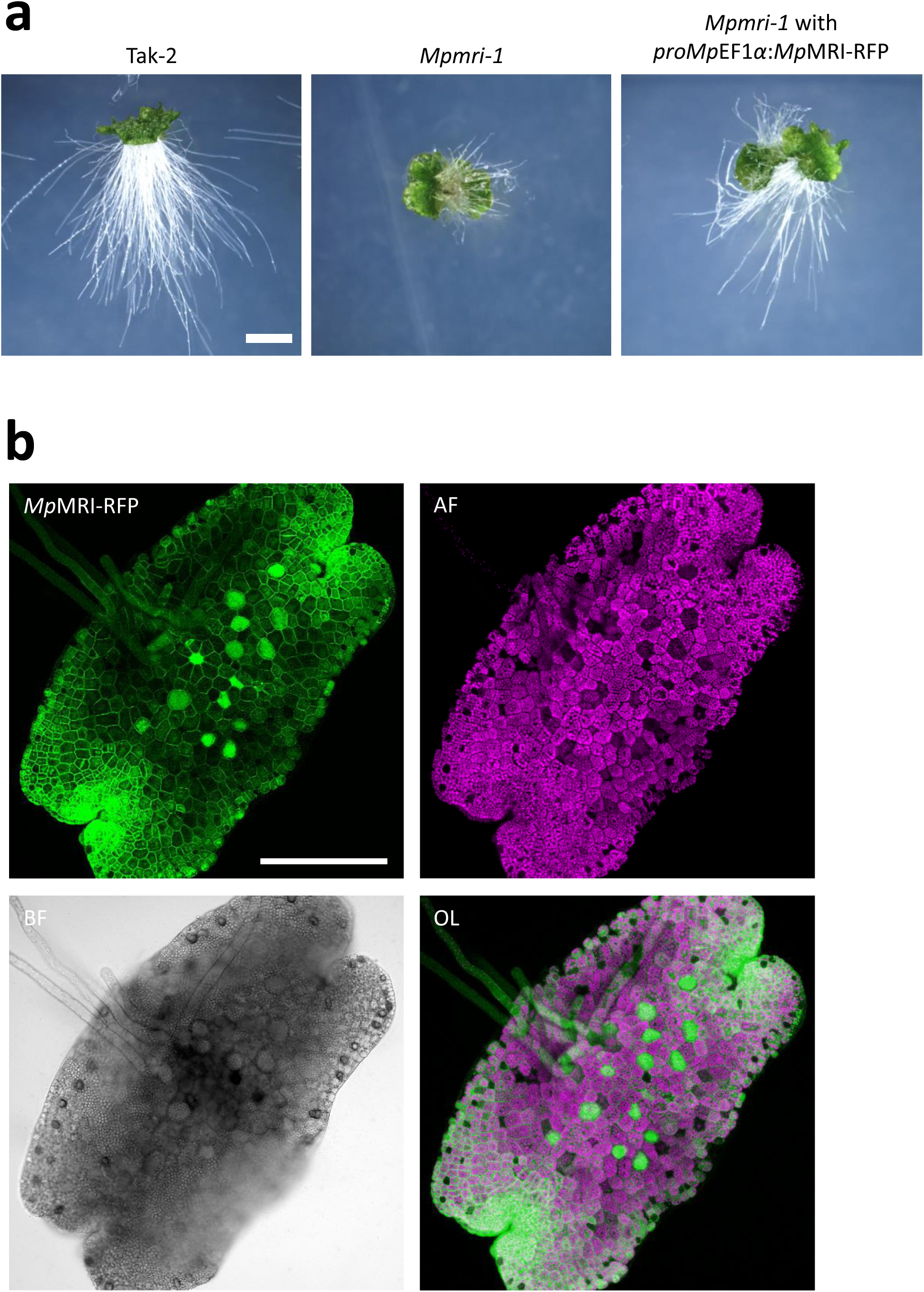
Restoration of rhizoid integrity and growth in *Mpmri-1* upon expression of *proMp*EF1α:*Mp*MRI-RFP. **(a)** Representative images of the data presented in Fig. 2a. Scale bar: 1 mm. **(b)** Maximum projection of a Z-stack of 1 day-old *Mpmri-1* gemmae expressing *proMp*EF1α:*Mp*MRI-RFP. Scale bar = 250 µm. RFP = red fluorescence protein. AF = autofluorescence. BF = bright field. OL = Overlay of RFP-derived fluorescence and autofluorescence channels.

**Supplementary Figure 3:**
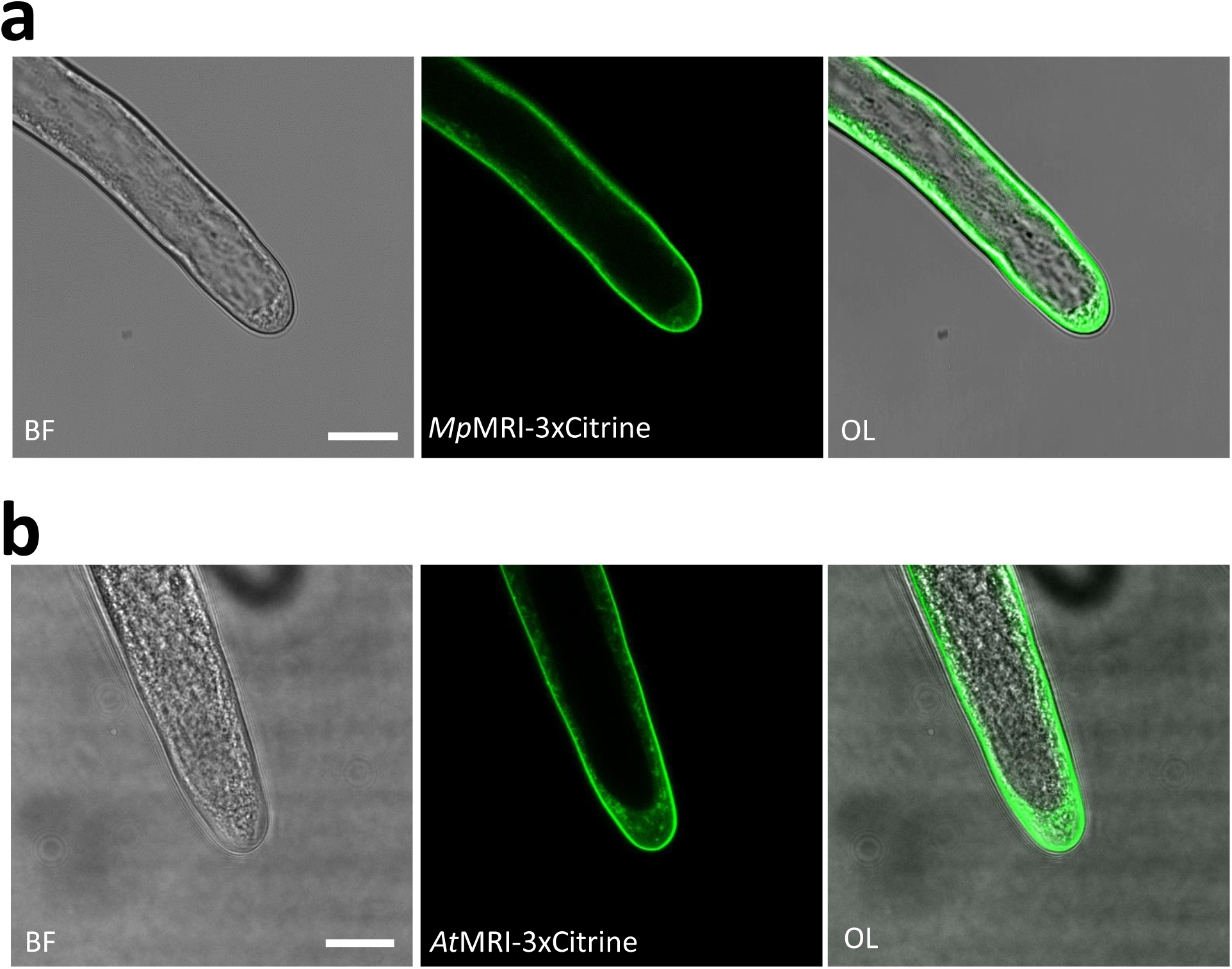
Localization of *Mp*MRI- and *At*MRI-fluorescent protein fusions in rhizoids. **(a)** *Mp*MRI-3xCitrine in Tak-2 x Tak-1 rhizoids. **(b)** *At*MRI-3xCitrine in Tak-2 x Tak-1 rhizoids. Scale bars = 20 µm. BF = bright field. OL = Overlay.

**Supplementary Figure 4:**
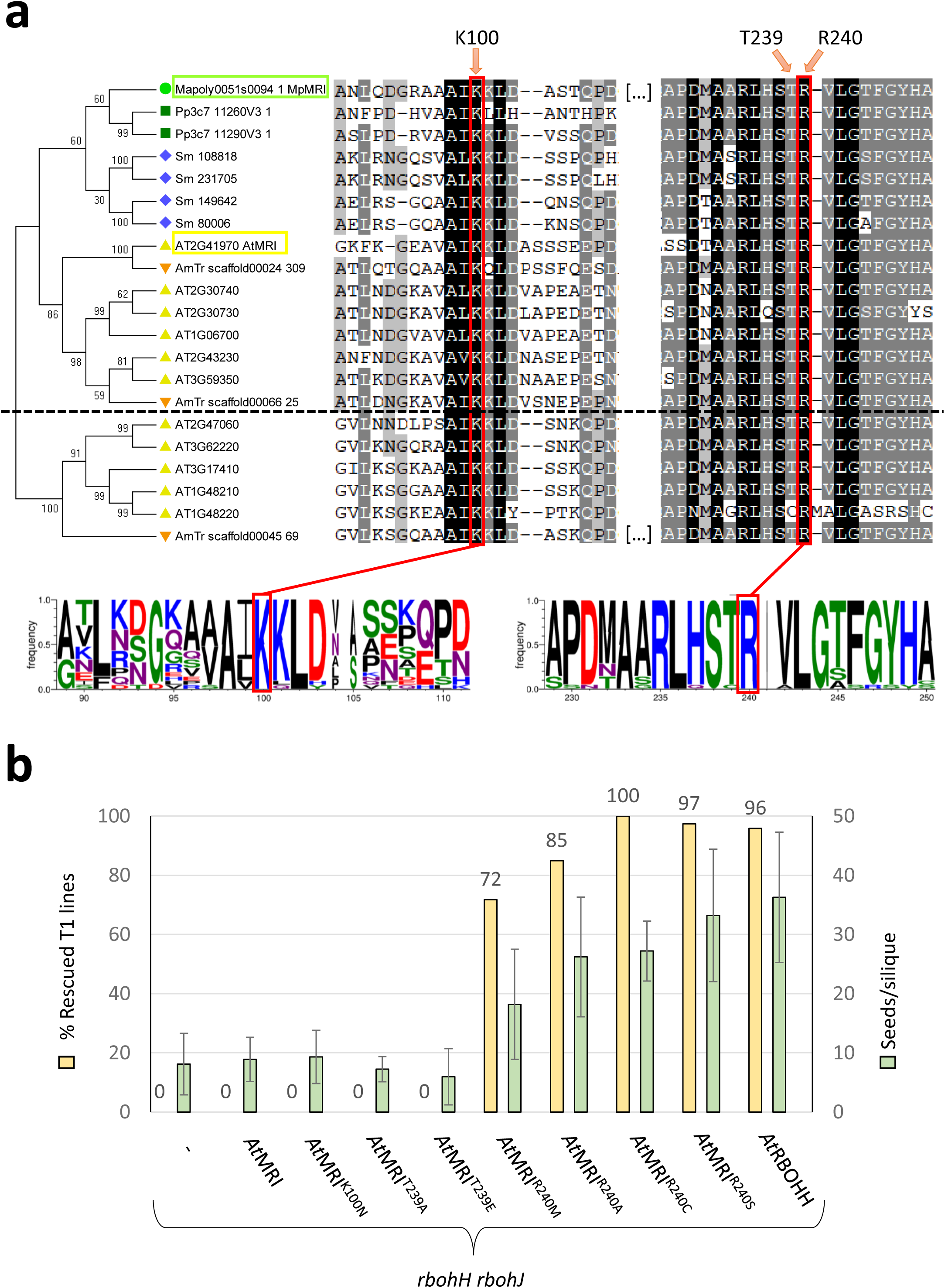
Structural and functional analysis of the STR-motif of PTI1-like receptor-like cytoplasmic kinases. **(a)** Phylogenetic analysis and multiple sequence alignment of land plant PTI1-like RLCK-VIII full-length amino acid sequences centered on the invariant K100 of kinase subdomain II and the conserved STR motif. Green circle, *M. polymorpha;* green squares, *Physcomitrella patens*; blue diamonds, *Selaginella moellendorffii*; orange triangles, *Amborella trichopoda*; yellow triangles, *A. thaliana*. Horizontal dashed line separates the ‘MRI-like’ subgroup (above) from the other PTI1-like RLCKs-VIII. Phylogenetic trees were calculated with MEGA7^53^ using the Maximum Likelihood method based on the JTT matrix-based model^54^ and 1000 repetitions. **(b)** Genetic rescue assay of *AtrbohH rbohJ* male sterility via pollen expression of *At*MRI structural variants. Shown is % of T1 lines that showed a sterility rescue, as well as the number of seeds per silique on average in the T1 lines. The *AtrbohH rbohJ* male sterility could only be rescued upon expression of either *At*RBOHH or any *At*MRI variants harboring a R240 substitution. Data for *AtrbohH rbohJ* sterility rescue by *AtRBOHH* expression is from Boisson-Dernier et al., 2013.

**Supplementary Figure 5:**
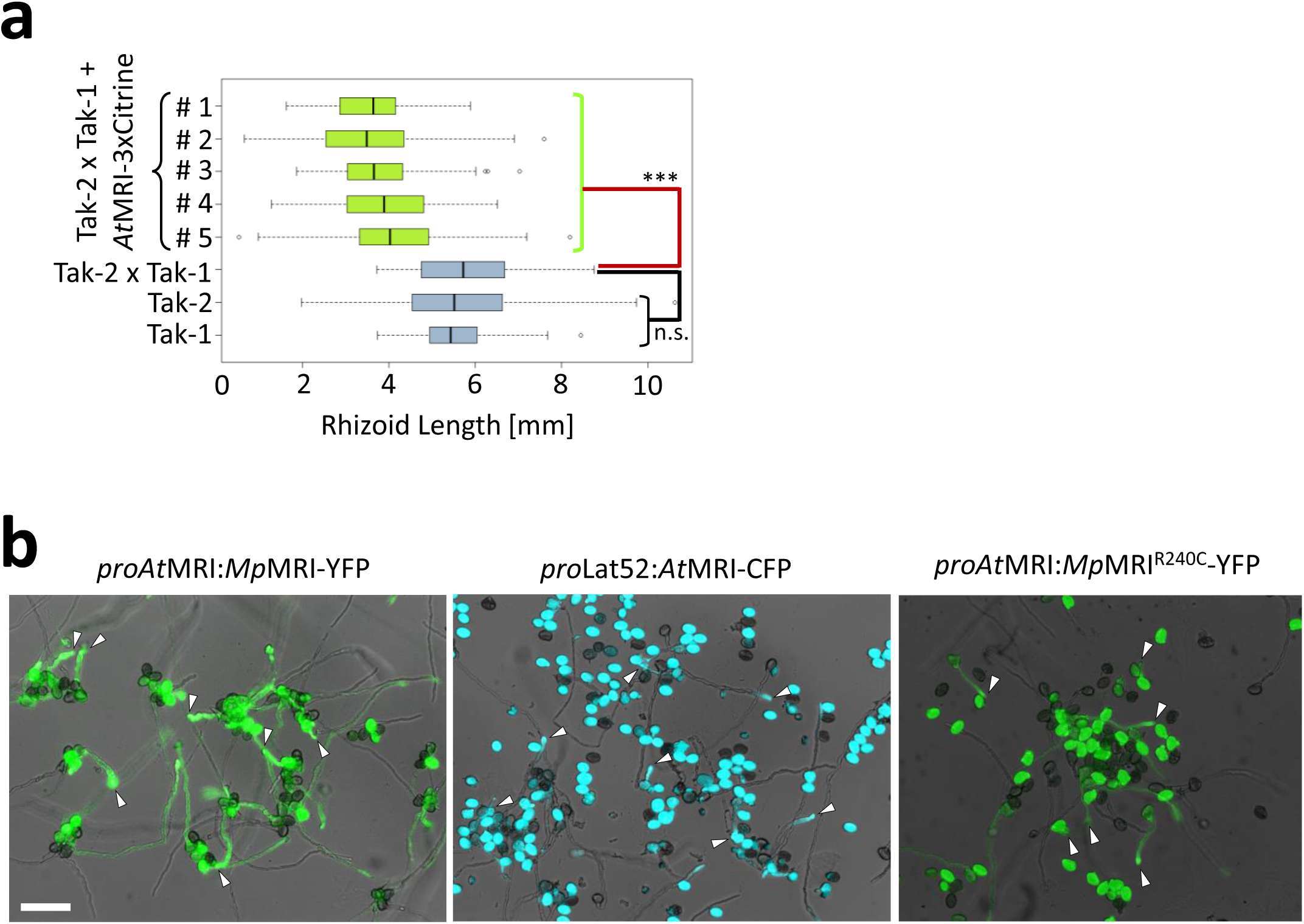
Pollen tube growth inhibition upon expression of *At*MRI, *Mp*MRI and *Mp*MRI^R240C^. **(a)** Rhizoid length of 7 d old Marchantia Tak-2 x Tak-1 wild-type thalli expressing *At*MRI compared to a non-transformed Tak-2 x Tak-1, Tak-2 and Tak-1 non-transformed controls (n > 10 thalli/independent line). Significance of all expression lines against non-transformed Tak-2 x Tak-1 was tested with a two-tailed, unpaired student’s T-test (***: p<0.001). Data of non-transformed line was statistically not significantly different (n.s.). **(b)** Representative overlay images for hemizygous lines whose data is shown in Fig. 5c, d and Supplementary Table 4. White arrowheads indicate short and occasionally wide pollen tubes. Scale bar: 80 µm.

**Supplementary Figure 6:**
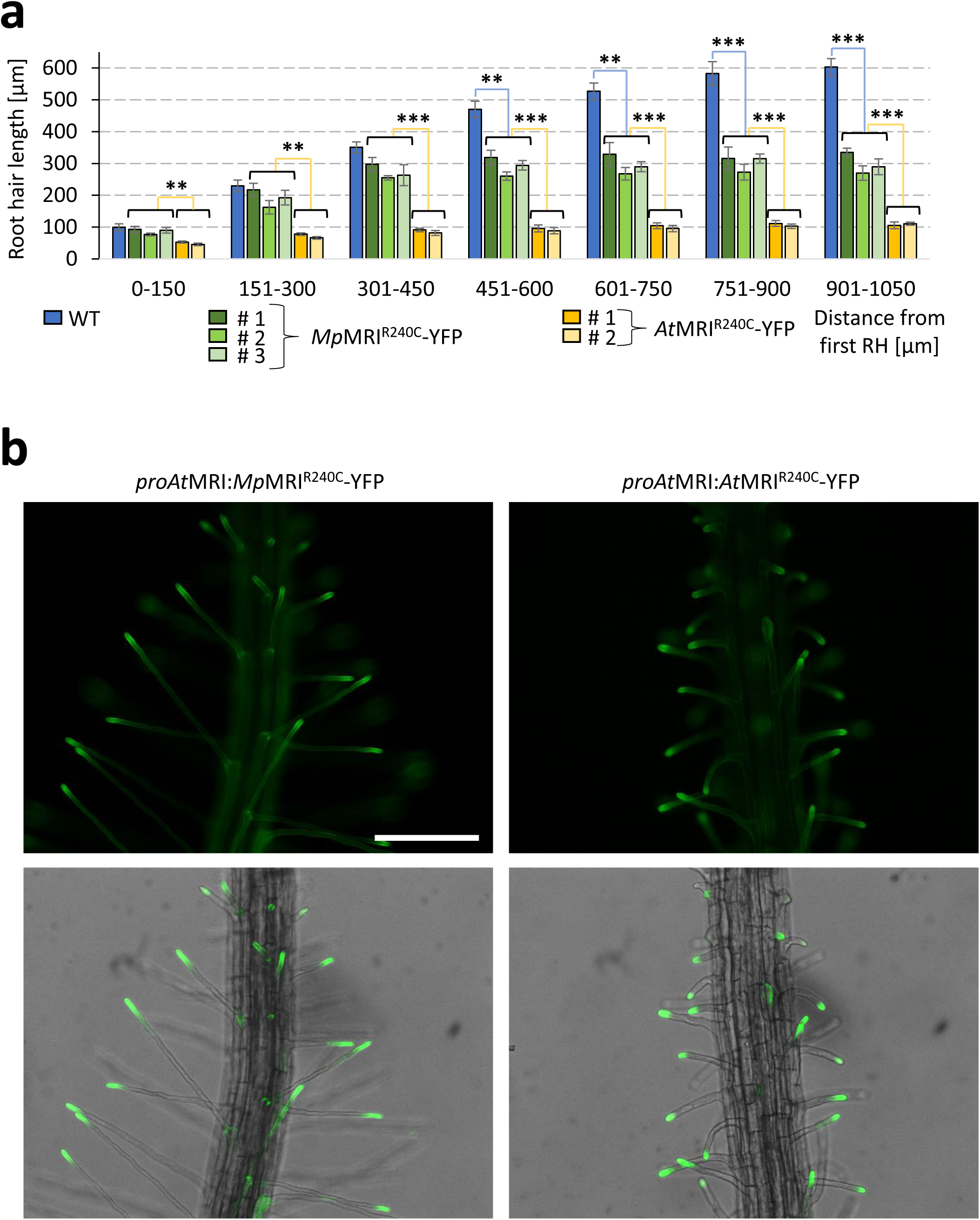
Expression of *At*MRI^R240C^-YFP and *Mp*MRI^R240C^-YFP causes growth inhibition in Arabidopsis root hairs. **(a)** Root hair length profile of wild-type seedlings and transgenic lines expressing *proAt*MRI:*Mp*MRI^R240C^-YFP (n = 3 independent lines) or *proAt*MRI:*At*MRI^R240C^-YFP (n = 2 independent lines). Each primary root was divided in optical sections of 150 μm and root hair length was determined for 5-10 root hairs per section. Shown are the mean values of n = 5 seedlings per genotype and section; error bars show the standard error from the mean. Significance tested with a two-tailed, unpaired student’s T-test; **: p<0.01, ***: p<0.001. **(b)** Fluorescence (top) and overlay (bottom) images of *proAt*MRI:*MpMR*I^R240C^-YFP and *proAt*MRI:*At*MRI^R240C^-YFP in root hairs, both localizing normally to the plasma membrane; scale bar: 200 µm.

**Supplementary Figure 7:**
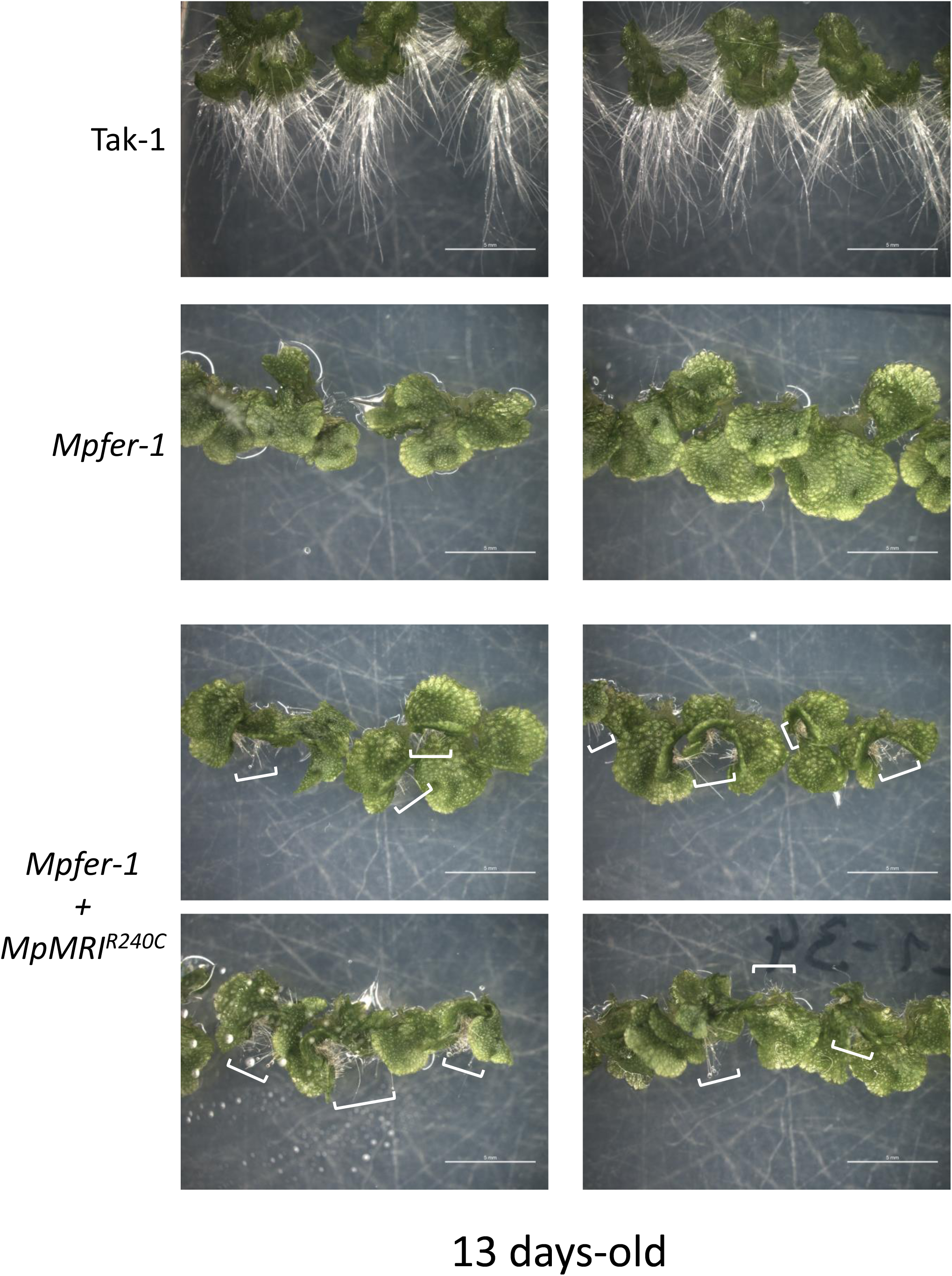
*Mp*MRI^R240C^-RFP expression partially rescues rhizoid bursting of *Mpfer-1* mutant. Representative images at 13 days of the lines analyzed in Fig. 6 sections (a), (b) and (c). White brackets depict the increase in rhizoid density in different *Mpfer-1* lines expressing *Mp*MRI^R240C^-RFP as compared to the rare rhizoids in untransformed *Mpfer-1*. Scale bars: 5 mm. See also Fig. 6 for younger stages.

## References

1. de Vries, J. & Archibald, J. M. Plant evolution: landmarks on the path to terrestrial life. New Phytol. 217, 1428–1434 (2018).

2. Rensing, S. A. Great moments in evolution: the conquest of land by plants. Curr. Opin. Plant Biol. 42, 49–54 (2018).

3. Bowman, J. L., Briginshaw, L. N. & Florent, S. N. Evolution and co-option of developmental regulatory networks in early land plants. Curr. Top. Dev. Biol. 131, 35– 53 (2019).

4. Jones, V. A. S. & Dolan, L. The evolution of root hairs and rhizoids. Ann. Bot. 110, 205–212 (2012).

5. Friedman, W. E. The evolutionary history of the seed plant male gametophyte. Trends Ecol. Evol. 8, 15–21 (1993).

6. Rounds, C. M. & Bezanilla, M. Growth Mechanisms in Tip-Growing Plant Cells. Annu. Rev. Plant Biol. 64, 243–265 (2013).

7. Franck, C. M., Westermann, J. & Boisson-Dernier, A. Plant Malectin-Like Receptor Kinases: From Cell Wall Integrity to Immunity and Beyond. Annu. Rev. Plant Biol. 69, 301–328 (2018).

8. Ge, Z., Cheung, A. Y. & Qu, L. Pollen tube integrity regulation in flowering plants: insights from molecular assemblies on the pollen tube surface. New Phytol. 222, 687– 693 (2019).

9. Wolf, S., Hématy, K. & Höfte, H. Growth control and cell wall signaling in plants. Annu. Rev. Plant Biol. 63, 381–407 (2012).

10. Engelsdorf, T. & Hamann, T. An update on receptor-like kinase involvement in the maintenance of plant cell wall integrity. Ann. Bot. 114, 1339–1347 (2014).

11. Hamann, T. The plant cell wall integrity maintenance mechanism – A case study of a cell wall plasma membrane signaling network. Phytochemistry 112, 100–109 (2015).

12. Boisson-Dernier, A. et al. Disruption of the pollen-expressed FERONIA homologs ANXUR1 and ANXUR2 triggers pollen tube discharge. Development 136, 3279–3288 (2009).

13. Miyazaki, S. et al. ANXUR1 and 2, sister genes to FERONIA/SIRENE, are male factors for coordinated fertilization. Curr. Biol. 19, 1327–31 (2009).

14. Ge, Z. et al. Arabidopsis pollen tube integrity and sperm release are regulated by RALF-mediated signaling. Science. 358, 1596–1600 (2017).

15. Mecchia, M. A. et al. RALF4/19 peptides interact with LRX proteins to control pollen tube growth in Arabidopsis. Science. 358, 1600–1603 (2017).

16. Boisson-Dernier, A. et al. ANXUR receptor-like kinases coordinate cell wall integrity with growth at the pollen tube tip via NADPH oxidases. PLoS Biol. 11, e1001719 (2013).

17. Franck, C. et al. The protein phosphatases ATUNIS1 and ATUNIS2 regulate cell wall integrity in tip-growing cells. Plant Cell 30, 1906–1923 (2018).

18. Boisson-Dernier, A., Franck, C. M., Lituiev, D. S. & Grossniklaus, U. Receptor-like cytoplasmic kinase MARIS functions downstream of CrRLK1L-dependent signaling during tip growth. Proc. Natl. Acad. Sci. U. S. A. 112, 12211–6 (2015).

19. Duan, Q., Kita, D., Li, C., Cheung, A. Y. & Wu, H.-M. FERONIA receptor-like kinase regulates RHO GTPase signaling of root hair development. Proc. Natl. Acad. Sci. U. S. A. 107, 17821–6 (2010).

20. Liao, H.-Z. et al. *MARIS* plays important roles in *Arabidopsis* pollen tube and root hair growth. J. Integr. Plant Biol. 58, 927–940 (2016).

21. Zhou, J., Loh, Y.-T., Bressan, R. A. & Martin, G. B. The tomato gene Pti1 encodes a serine/threonine kinase that is phosphorylated by Pto and is involved in the hypersensitive response. Cell 83, 925–935 (1995).

22. Zhang, L. et al. CARK1 mediates ABA signaling by phosphorylation of ABA receptors. Cell Discov. 4, 30 (2018).

23. Honkanen, S. et al. The Mechanism Forming the Cell Surface of Tip-Growing Rooting Cells Is Conserved among Land Plants. Curr. Biol. 26, 3238–3244 (2016).

24. Bowman, J. L. et al. Insights into Land Plant Evolution Garnered from the Marchantia polymorpha Genome. Cell 171, 287–304 (2017).

25. Sessa, G., D’Ascenzo, M. & Martin, G. B. The major site of the Pti1 kinase phosphorylated by the Pto kinase is located in the activation domain and is required for Pto-Pti1 physical interaction. Eur. J. Biochem. 267, 171–178 (2000).

26. Anthony, R. G., Khan, S., Costa, J., Pais, M. S. & Bögre, L. The Arabidopsis protein kinase PTI1-2 is activated by convergent phosphatidic acid and oxidative stress signaling pathways downstream of PDK1 and OXI1. J. Biol. Chem. 281, 37536–46 (2006).

27. Herrmann, M. M., Pinto, S., Kluth, J., Wienand, U. & Lorbiecke, R. The PTI1-like kinase ZmPti1a from maize (Zea mays L.) co-localizes with callose at the plasma membrane of pollen and facilitates a competitive advantage to the male gametophyte. BMC Plant Biol. 6, 22 (2006).

28. Takahashi, A. et al. Rice Pti1a Negatively Regulates RAR1-Dependent Defense Responses. Plant Cell 19, 2940–2951 (2007).

29. Twell, D., Yamaguchi, J., Wing, R. A., Ushiba, J. & McCormick, S. Promoter analysis of genes that are coordinately expressed during pollen development reveals pollen-specific enhancer sequences and shared regulatory elements. Genes Dev. 5, 496– 507 (1991).

30. Matsui, H., Yamazaki, M., Kishi-Kaboshi, M., Takahashi, A. & Hirochika, H. AGC Kinase OsOxi1 Positively Regulates Basal Resistance through Suppression of OsPti1a-Mediated Negative Regulation. Plant Cell Physiol. 51, 1731–1744 (2010).

31. Boisson-Dernier, A., Franck, C. M., Lituiev, D. S. & Grossniklaus, U. Receptor-like cytoplasmic kinase MARIS functions downstream of *Cr* RLK1L-dependent signaling during tip growth. Proc. Natl. Acad. Sci. U. S. A. 112, 12211–12216 (2015).

32. Carol, R. J. & Dolan, L. Building a hair: tip growth in Arabidopsis thaliana root hairs. Philos. Trans. R. Soc. Lond. B. Biol. Sci. 357, 815–21 (2002).

33. Menand, B. et al. An Ancient Mechanism Controls the Development of Cells with a Rooting Function in Land Plants. Science. 316, 1477–1480 (2007).

34. Proust, H. et al. RSL Class I Genes Controlled the Development of Epidermal Structures in the Common Ancestor of Land Plants. Curr. Biol. 26, 93–99 (2016).

35. Scotland, R. W. Deep homology: A view from systematics. BioEssays 32, 438–449 (2010).

36. Hackenberg, D. & Twell, D. The evolution and patterning of male gametophyte development. Curr. Top. Dev. Biol. 131, 257–298 (2019).

37. Doyle, J. A. Seed ferns and the origin of angiosperms. J. Torrey Bot. Soc. 133, 169– 209 (2006).

38. Williams, J. H. Novelties of the flowering plant pollen tube underlie diversification of a key life history stage. Proc. Natl. Acad. Sci. U. S. A. 105, 11259–11263 (2008).

39. Liao, H.-Z. et al. *MARIS* plays important roles in *Arabidopsis* pollen tube and root hair growth. J. Integr. Plant Biol. 58, 927–940 (2016).

40. Schwizer, S. et al. The Tomato Kinase Pti1 Contributes to Production of Reactive Oxygen Species in Response to Two Flagellin-Derived Peptides and Promotes Resistance to *Pseudomonas syringae* Infection. Mol. Plant-Microbe Interact. 30, 725– 738 (2017).

41. Oh, S.-K. et al. Cucumber Pti1-L is a cytoplasmic protein kinase involved in defense responses and salt tolerance. J. Plant Physiol. 171, 817–822 (2014).

42. Matsui, H. et al. Plasma Membrane Localization Is Essential for *Oryza sativa* Pto-Interacting Protein 1a-Mediated Negative Regulation of Immune Signaling in Rice. Plant Physiol. 166, 327–336 (2014).

43. Tougane, K. et al. Evolutionarily conserved regulatory mechanisms of abscisic acid signaling in land plants: characterization of ABSCISIC ACID INSENSITIVE1-like type 2C protein phosphatase in the liverwort Marchantia polymorpha. Plant Physiol. 152, 1529–43 (2010).

44. Jahan, A. et al. Archetypal Roles of an Abscisic Acid Receptor in Drought and Sugar Responses in Liverworts. Plant Physiol. 179, 317–328 (2019).

45. Eklund, D. M. et al. An Evolutionarily Conserved Abscisic Acid Signaling Pathway Regulates Dormancy in the Liverwort Marchantia polymorpha. Curr. Biol. 28, 3691–3699.e3 (2018).

46. Carella, P., Gogleva, A., Tomaselli, M., Alfs, C. & Schornack, S. Phytophthora palmivora establishes tissue-specific intracellular infection structures in the earliest divergent land plant lineage. Proc. Natl. Acad. Sci. U. S. A. 115, E3846–E3855 (2018).

47. Clough, S. J. & Bent, A. F. Floral dip: a simplified method for Agrobacterium-mediated transformation of Arabidopsis thaliana. Plant J. 16, 735–43 (1998).

48. Ishizaki, K., Chiyoda, S., Yamato, K. T. & Kohchi, T. Agrobacterium-Mediated Transformation of the Haploid Liverwort Marchantia polymorpha L., an Emerging Model for Plant Biology. Plant Cell Physiol. 49, 1084–1091 (2008).

49. Kubota, A., Ishizaki, K., Hosaka, M. & Kohchi, T. Efficient *Agrobacterium* - Mediated Transformation of the Liverwort *Marchantia polymorpha* Using Regenerating Thalli. Biosci. Biotechnol. Biochem. 77, 167–172 (2013).

50. Johnson, C. M., Stout, P. R., Broyer, T. C., & Carlton, A. B. Comparative chlorine requirements of different plant species. Plant Soil 8, 337–358 (1957).

51. Boavida, L. C. & McCormick, S. Temperature as a determinant factor for increased and reproducible in vitro pollen germination in Arabidopsis thaliana. Plant J. 52, 570– 82 (2007).

52. Larkin, M. A. et al. Clustal W and Clustal X version 2.0. Bioinformatics 23, 2947–8 (2007).

53. Kumar, S., Stecher, G. & Tamura, K. MEGA7: Molecular Evolutionary Genetics Analysis Version 7.0 for Bigger Datasets. Mol. Biol. Evol. 33, 1870–1874 (2016).

54. Jones, D. T., Taylor, W. R. & Thornton, J. M. The rapid generation of mutation data matrices from protein sequences. Comput. Appl. Biosci. 8, 275–82 (1992).

55. Schindelin, J. et al. Fiji: an open-source platform for biological-image analysis. Nat. Methods 9, 676–682 (2012).

