## Supplementary Tables for "An Evolutionarily Conserved Receptor-like Kinases Signaling Module Controls Cell Wall Integrity During Tip-Growth"

**Supplementary Table 1: The 10 closest *Mp*MRI homologs of *Arabidopsis thaliana*.** Shown is the output of a pBlast using the *Mp*MRI (*Mapoly0051s0094*) amino acid sequence as query against the Araport11 database (carried out on https://www.arabidopsis.org/Blast/ using standard settings). (a) Comparison of full-length amino acid sequences; (b) Comparison of kinase domain only.

| (a) **Full-length amino acid sequences** | | | |
|---|---|---|---|
| **Gene** | **% identity (region spanning)** | **% similarity (region spanning)** | **E-value** |
| AT2G30740 | 70 (254/358) | 79 (286/358) | e-147 |
| AT3G59350 | 69 (253/364) | 79 (290/364) | e-145 |
| AT2G43230 | 69 (250/361) | 79 (288/361) | e-144 |
| AT2G47060 | 69 (254/363) | 77 (283/363) | e-143 |
| AT1G06700 | 68 (248/360) | 77 (280/360) | e-141 |
| AT3G17410 | 73 (248/338) | 81 (274/338) | e-141 |
| AT3G62220 | 67 (246/366) | 78 (289/366) | e-141 |
| AT1G48210 | 67 (245/365) | 76 (280/365) | e-138 |
| AT2G30730 | 69 (225/323) | 82 (265/323) | e-133 |
| **AT2G41970 (*At*MRI)** | 64 (233/364) | 76 (278/364) | e-133 |
| (b) **Kinase domain amino acid sequences** | | | |
| **Gene** | **% identity (region spanning)** | **% similarity (region spanning)** | **E-value** |
| AT2G43230 | 81 (230/283) | 89 (254/283) | e-137 |
| AT3G59350 | 80 (229/283) | 89 (253/283) | e-137 |
| AT2G30740 | 81 (230/283) | 88 (251/283) | e-137 |
| AT1G06700 | 81 (230/283) | 88 (251/283) | e-136 |
| AT3G17410 | 81 (229/281) | 87 (247/281) | e-134 |
| AT2G47060 | 79 (223/281) | 86 (242/281) | e-130 |
| AT3G62220 | 77 (218/281) | 86 (244/281) | e-129 |
| AT1G48210 | 77 (218/281) | 85 (240/281) | e-128 |
| AT2G30730 | 74 (212/283) | 85 (241/283) | e-126 |
| **AT2G41970 (*At*MRI)** | 73 (209/283) | 86 (244/283) | e-124 |

**Supplementary Table 2: Segregation analysis of the *mri-1* allele in the *mri-1*/MRI background transformed with *proAt*MRI:*Mp*MRI-YFP.** Shown are the observed numbers of T2 progeny - which were wild-type, heterozygous or homozygous for the *mri-1*-allele. Observed values of the transformed lines were compared by means of a chi-square test of independence with the expected values for a mendelian distribution or a male gametophytic transmission defect. [*] marks data from Boisson-Dernier et al., 2015.

| | **MRI/MRI** | ***mri-1*/MRI** | ***mri-1/mri-1*** | **n** | **Ratio** | **p-value (two-tailed Chi-square test)** |
|---|---|---|---|---|---|---|
| Observed (*mri-1*/MRI) [*] | 183 | 170 | 0 | 353 | 1 : 0.93 : 0 | |
| Observed (*mri-1*/MRI with *At*MRI-YFP) [*] | 23 | 38 | 29 | 90 | 1 : 1.65 : 1.26 | |
| Observed (*mri-1*/MRI with *Mp*MRI-YFP) | 69 | 124 | 7 | 200 | 1 : 1.80 : 0.10 | |
| Expected (Mendelian distribution) | 50 | 100 | 50 | 200 | 1 : 2 : 1 | <0.001 |
| Expected (Male gametophytic defect) | 100 | 100 | 0 | 200 | 1 : 1 : 0 | <0.001 |

**Supplementary Table 3:** Structural *At*MRI variants and the effect of their expression on male transmission of the *mri-1* allele in the progeny of heterozygous Arabidopsis *mri-1*/MRI plants. All fusion proteins are under the control of the pollen specific promoter *proLat52*. Grey, negative control. Green, positive control. Blue, overactive variant. Orange, variants related to K100. Yellow, variants related to T239. [*] marks data from Boisson-Dernier et al., 2015.

| **Transgene** | **MRI/MRI** | ***mri-1*/MRI** | ***mri-1/mri-1*** |
|---|---|---|---|
| *At*ANX1-YFP | 47 | 49 | 0 |
| *At*MRI-YFP [*] | 23 | 38 | 29 |
| *At*MRI$^{R240C}$-YFP | 28 | 43 | 17 |
| *At*MRI$^{K100N}$-CFP | 23 | 49 | 13 |
| *At*MRI$^{K100E}$-CFP | 29 | 50 | 12 |
| *At*MRI$^{T239A}$-CFP | 31 | 47 | 18 |
| *At*MRI$^{T239E}$-CFP | 25 | 47 | 20 |

**Supplementary Table 4: Pollen tube growth inhibition upon expression of *At*MRI, *Mp*MRI and their R240C structural variants.** Shown is the effect of hemizygous transgene expression on the percentage of fluorescent pollen grains and tubes. Each entry represents an independent transgenic line.

| Transgene | | Fluorescent pollen grains [%] | n | Fluorescent pollen tubes [%] | n |
|---|---|---|---|---|---|
| *proAt*MRI:*Mp*MRI-YFP (hemiz.) | # 1 | 48 | 104 | 55 | 698 |
| | # 2 | 48 | 71 | 54 | 693 |
| | # 3 | 50 | 64 | 48 | 728 |
| *proAt*MRI:*Mp*MRI$^{R240C}$-YFP (hemiz.) | # 1 | 52 | 386 | 43 | 317 |
| | # 2 | 56 | 424 | 41 | 250 |
| | # 3 | 53 | 489 | 37 | 412 |
| *pro*Lat52:*At*MRI-CFP (hemiz.) | | 58 | 110 | 16 | 438 |
| *pro*Lat52:*At*MRI$^{R240C}$-CFP (hemiz.) | | 39 | 337 | 0 | 250 |

**Supplementary Table 5: Genetically modified *Arabidopsis thaliana* lines.** Lines are separated into mutant and transgenic lines. (a) Mutant lines, including targeted genetic locus, plant ecotype and molecular markers. (b) Transgenic lines, including transformed construct, genotyping primers and plant selection markers.

| **(a) Mutant lines** | **Targeted locus** | **Ecotype** | **Molecular markers** |
|---|---|---|---|
| *mri-1*/MRI (*qrt*), (CSHL_GT21229) | AT2G41970 | Ler originally but outcrossed 4 times with Col-0 (Boisson-Dernier et al., 2015). | PCR (MRI): ABD681/ABD682 |
| | | | PCR (*mri-1*): ABD682/ABD747 |
| | | | Selection *mri-1*: Kanamycin |
| *rbohH-3 rbohJ-3,* (SALK_136917; SALK_050665) | AT5G60010<br>AT3G45810 | Col-0 | See Boisson-Dernier et al., 2013 |
| **(b) Transgenic lines** | **Transformed construct** | **Genotyping primers** | **Plant selection markers** |
| Col-0 with *proAt*MRI:*At*MRI$^{R240C}$-YFP | pABD85 | Construct: ABD766/ABD736 | Construct: Basta |
| Col-0 with *proAt*MRI:*Mp*MRI-YFP | pJW20 | Construct: ABD766/JW35 | Construct: Basta |
| Col-0 with *proAt*MRI:*Mp*MRI$^{R240C}$-YFP | pJW36 | Construct: ABD766/JW35 | Construct: Basta |
| Col-0 with *pro*Lat52:*At*MRI$^{R240C}$-CFP | pABD49 | Construct: ABD766/JW35 | Construct: Basta |
| *mri-1* (*qrt*) with *pro*Lat52:*At*MRI$^{R240C}$-CFP | pABD49 | *mri-1*: ABD682/ABD747 | Mutant: Kanamycin |
| | | Construct: ABD766/JW35 | Construct: Basta |
| *mri-1* (*qrt*) with *proAt*MRI:*At*MRI-YFP | pABD84 | *mri-1*: ABD682/ABD747 | Mutant: Kanamycin |
| | | Construct: ABD766/ABD736 | Construct: Basta |
| *mri-1* (*qrt*) with *proAt*MRI:*Mp*MRI-YFP | pJW20 | *mri-1*: ABD682/ABD747 | Mutant: Kanamycin |
| | | Construct: ABD766/JW35 | Construct: Basta |
| *mri-1* (*qrt*) with *pro*Lat52:*At*ANX1-YFP | pABD39 | *mri-1*: ABD682/ABD747 | Mutant: Kanamycin |
| | | Construct: ABD546/ABD630 | Construct: Basta |
| *mri-1* (*qrt*) or *rbohH-3 rbohJ-3* with *pro*Lat52:*At*MRI-YFP | pABD46 | Construct: ABD680/ABD630 | Construct: Basta |
| *mri-1* (*qrt*) or *rbohH-3 rbohJ-3* with *pro*Lat52:*At*MRI$^{R240C}$-YFP | pABD48 | Construct: ABD680/ABD630 | Construct: Basta |
| *mri-1* (*qrt*) or *rbohH-3 rbohJ-3* with *pro*Lat52:*At*MRI$^{K100N}$-YFP | pABD88 | Construct: ABD680/ABD630 | Construct: Basta |
| *mri-1* (*qrt*) or *rbohH-3 rbohJ-3* with *pro*Lat52:*At*MRI$^{K100E}$-YFP | pABD108 | Construct: ABD680/ABD630 | Construct: Basta |
| *mri-1* (*qrt*) or *rbohH-3 rbohJ-3* with *pro*Lat52:*At*MRI$^{T239A}$-YFP | pABD89 | Construct: ABD680/ABD630 | Construct: Basta |
| *mri-1* (*qrt*) or *rbohH-3 rbohJ-3* with *pro*Lat52:*At*MRI$^{T239E}$-YFP | pABD90 | Construct: ABD680/ABD630 | Construct: Basta |
| *mri-1* (*qrt*) or *rbohH-3 rbohJ-3* with *pro*Lat52:*At*MRI$^{R240M}$-YFP | pABD109 | Construct: ABD680/ABD630 | Construct: Basta |
| *mri-1* (*qrt*) or *rbohH-3 rbohJ-3* with *pro*Lat52:*At*MRI$^{R240A}$-YFP | pABD110 | Construct: ABD680/ABD630 | Construct: Basta |
| *mri-1* (*qrt*) or *rbohH-3 rbohJ-3* with *pro*Lat52:*At*MRI$^{R240S}$-YFP | pABD111 | Construct: ABD680/ABD630 | Construct: Basta |
| *rbohH-3 rbohJ-3* with *proAt*ACA9-GFP-RbohH | pABD18 | See Boisson-Dernier et al., 2013 | Construct: Hygromycin |

**Supplementary Table 6: Genetically modified *Marchantia polymorpha* lines.** Lines are separated into mutant and transgenic lines. (a) Mutant lines, including targeted genetic locus, plant ecotype and molecular markers. (b) Transgenic lines, including transformed construct, genotyping primers and plant selection markers.

| (a) Mutant lines | Targeted locus | Ecotype | Molecular markers |
|---|---|---|---|
| *Mpmri-1*<br><br>(Honkanen et al., 2016; [1]) | *MpMRI* | Tak-2 | PCR (*Mpmri-1*): JW60/JW63<br><br>Selection (T-DNA): Hygromycin |
| *Mpfer-1*<br><br>(Honkanen et al., 2016; [1]) | *MpFER* | Tak-1 | PCR (*Mp*FER): JW52/JW59<br><br>PCR (*Mpfer-1*): JW52/JW64<br><br>Selection (T-DNA): Hygromycin |
| **(b) Transgenic lines** | **Transgene** | **Genotyping primers** | **Plant selection markers** |
| *Mpmri-1* with<br>*proMp*EF1α:*At*MRI-RFP | pJW10 | *Mpmri-1*: JW60/JW63<br>Construct: JW17/ABD736 | Construct: Chlorsulfuron<br>Mutant: Hygromycin |
| *Mpmri-1* with<br>*proMp*EF1α:*Mp*MRI-RFP | pJW28 | *Mpmri-1*: JW60/JW63<br>Construct: JW17/JW35 | Construct: Chlorsulfuron<br>Mutant: Hygromycin |
| *Mpfer-1* (or Tak-2 x Tak-1) with<br>*proMp*EF1α:*Mp*MRI$^{R240C}$-RFP | pJW51 | *Mpfer-1*: JW52/JW64<br>Construct: JW17/JW35 | Construct: Chlorsulfuron<br>Mutant: Hygromycin |
| Tak-2 x Tak-1 with<br>*proMp*EF1α:*At*MRI-3xCitrine | pJW09 | Construct: JW17/ABD736 | Construct: (Gentamicin) |
| Tak-2 x Tak-1 with<br>*proMp*EF1α:*Mp*MRI-3xCitrine | pJW27 | Construct: JW17/JW35 | Construct: (Gentamicin) |

**Supplementary Table 7: Vectors used in this study.** Vectors are separated into Gateway entry vectors/clones (a), destination vectors (b) and final expression constructs (c).

(a) Gateway entry vectors and clones

| Vector ID | Full name | Purpose | Bacterial selection marker | Vector type |
|---|---|---|---|---|
| pABD20 | *At*ANX1 (without stop) in pDONR221 | Cloning of *At*ANX1 ORF | Kanamycin | Entry clone |
| pABD43 | *At*MRI (without stop) in pDONR207 | Cloning of *At*MRI ORF | Gentamicin | Entry clone |
| pABD44 | *At*MRI$^{R240C}$ (without stop) in pDONR207 | Cloning of *At*MRI$^{R240C}$ | Gentamicin | Entry clone |
| pABD74 | *At*MRI$^{K100N}$ (without stop) in pDONR207 | Cloning of *At*MRIK$^{100N}$ | Gentamicin | Entry clone |
| pABD75 | *At*MRI$^{T239A}$ (without stop) in pDONR207 | Cloning of *At*MRI$^{T239A}$ | Gentamicin | Entry clone |
| pABD76 | *At*MRI$^{T239E}$ (without stop) in pDONR207 | Cloning of *At*MRI$^{T239E}$ | Gentamicin | Entry clone |
| pABD100 | *At*MRI$^{K100E}$ (without stop) in pDONR207 | Cloning of *At*MRI$^{K100E}$ | Gentamicin | Entry clone |
| pABD101 | *At*MRI$^{R240A}$ (without stop) in pDONR207 | Cloning of *At*MRI$^{R240A}$ | Gentamicin | Entry clone |
| pABD102 | *At*MRI$^{R240M}$ (without stop) in pDONR207 | Cloning of *At*MRI$^{R240M}$ | Gentamicin | Entry clone |
| pABD103 | *At*MRI$^{R240S}$ (without stop) in pDONR207 | Cloning of *At*MRI$^{R240S}$ | Gentamicin | Entry clone |
| pJW18 | *Mp*MRI (without stop) in pDONR207 | Cloning of *Mp*MRI | Gentamicin | Entry clone |
| pJW35 | *Mp*MRI$^{R240C}$ (without stop) in pDONR207 | Cloning of *Mp*MRI$^{R240C}$ | Gentamicin | Entry clone |

(b) Gateway destination vectors

| Vector ID | Full name | Purpose | Bacterial selection marker | Plant selection marker |
|---|---|---|---|---|
| pABD34 | *pro*Lat52:GW-YFP | Pollen-specific expression in *Arabidopsis* | Spectinomycin | Basta |
| pABD35 | *pro*Lat52:GW-CFP | Pollen-specific expression in *Arabidopsis* | Spectinomycin | Basta |
| pABD83 | *proAt*MRI:GW-YFP | pollen- and root hair-specific expression in *Arabidopsis* | Spectinomycin | Basta |
| pABD106 | *pro*MpEF1α:GW-3xCitrine (pMpGWB224; Ishizaki et al., 2015) | ubiquitous expression in *Marchantia* | Spectinomycin | Gentamicin |
| pABD107 | *pro*MpEF1α:GW-RFP (pMpGWB327; Ishizaki et al., 2015) | ubiquitous expression in *Marchantia* | Spectinomycin | Chlorsulfuron |

(c) Binary and Gateway expression vectors

| Vector ID | Full name | Entry clone and destination vector | Bacterial selection marker | Plant selection marker |
|---|---|---|---|---|
| pABD18 | *proAt*ACA9-GFP-RbohH | n/a | Kanamycin | Hygromycin |
| pABD39 | *pro*Lat52:*At*ANX1-YFP | pABD20/pABD34 | Spectinomycin | Basta |
| pABD46 | *pro*Lat52:*At*MRI-YFP | pABD43/pABD34 | Spectinomycin | Basta |

| | | | | |
|---|---|---|---|---|
| pABD47 | *pro*Lat52:*At*MRI-CFP | pABD43/pABD35 | Spectinomycin | Basta |
| pABD48 | *pro*Lat52:*At*MRI$^{R240C}$-YFP | pABD44/pABD34 | Spectinomycin | Basta |
| pABD49 | *pro*Lat52:*At*MRI$^{R240C}$-CFP | pABD44/pABD35 | Spectinomycin | Basta |
| pABD84 | *proAt*MRI:*At*MRI-YFP | pABD43/pABD83 | Spectinomycin | Basta |
| pABD85 | *proAt*MRI:*At*MRI$^{R240C}$-YFP | pABD44/pABD83 | Spectinomycin | Basta |
| pABD88 | *pro*Lat52:*At*MRI$^{K100N}$-CFP | pABD74/pABD35 | Spectinomycin | Basta |
| pABD89 | *pro*Lat52:*At*MRI$^{T239A}$-CFP | pABD75/pABD35 | Spectinomycin | Basta |
| pABD90 | *pro*Lat52:*At*MRI$^{T239E}$-CFP | pABD76/pABD35 | Spectinomycin | Basta |
| pABD108 | *pro*Lat52:*At*MRI$^{K100E}$-CFP | pABD100/pABD35 | Spectinomycin | Basta |
| pABD109 | *pro*Lat52:*At*MRI$^{R240A}$-CFP | pABD101/pABD35 | Spectinomycin | Basta |
| pABD110 | *pro*Lat52:*At*MRI$^{R240M}$-CFP | pABD102/pABD35 | Spectinomycin | Basta |
| pABD111 | *pro*Lat52:*At*MRI$^{R240S}$-CFP | pABD103/pABD35 | Spectinomycin | Basta |
| pJW09 | *proMp*EF1α:*At*MRI-3xCitrine | pABD43/pABD106 | Spectinomycin | Gentamicin |
| pJW10 | *proMp*EF1α:*At*MRI-RFP | pABD43/pABD107 | Spectinomycin | Chlorsulfuron |
| pJW20 | pro*At*MRI:*Mp*MRI-YFP | pJW18/pABD83 | Spectinomycin | Basta |
| pJW27 | *proMp*EF1α:*Mp*MRI-3xCitrine | pJW18/pABD106 | Spectinomycin | Gentamicin |
| pJW28 | *proMp*EF1α:*Mp*MRI-RFP | pJW18/pABD107 | Spectinomycin | Chlorsulfuron |
| pJW36 | *proAt*MRI:*Mp*MRI$^{R240C}$-YFP | pJW35/pABD83 | Spectinomycin | Basta |
| pJW51 | *proMp*EF1α:*Mp*MRI$^{R240C}$-RFP | pJW35/pABD107 | Spectinomycin | Chlorsulfuron |

### Supplementary Table 8: Oligonucleotides used in this study.

| Primer ID | Full name | Nucleotide Sequence (5' to 3') | Purpose |
|---|---|---|---|
| ABD546 | ANX1-F5 | GAAATTTGCAGACACGGCGGAG | Genotyping of *At*ANX1 ORF |
| ABD630 | eYFP-R | AAGCACTGCAGGCCGTAGC | Genotyping of ORF-YFP fusions |
| ABD681 | GT21229-F1 | TTCGGCTACCACGCTCCAGA | Genotyping of MRI/*mri-1* |
| ABD682 | GT21229-R1 | GGACCGGCCGGTTTAGAGTT | Genotyping of MRI/*mri-1* |
| ABD698 | mCFP-F | CGAGGAGCTGTTCACCGGGG | Genotyping of CFP- or YFP-ORF fusions |
| ABD699 | mCFP-R | CCTCGAACTTCACCTCGGCGC | Genotyping of ORF-CFP fusions |
| ABD747 | Ds5-2 | TCCGTTCCGTTTTCGTTTTTTAC | Genotyping of *mri-1*/MRI |
| ABD766 | p*At*MRI-F | TTGTTTCCCGACTTTATCGCGG | Genotyping of *proAt*MRI-ORF fusions |
| ABD756 | MRI_K100N_F | GGAGAAGCTGTTGCTATCAATAAACTTGATGCTAGTTCTTC | Site-directed mutagenesis |
| ABD757 | MRI_K100N_R | GAAGAACTAGCATCAAGTTTATTGATAGCAACAGCTTCTCC | Site-directed mutagenesis |
| ABD758 | MRI_T239A_F | GGCTAGGCTTCATTCTGCTCGTGTTTTGGGAAC | Site-directed mutagenesis |
| ABD759 | MRI_T239A_R | GTTCCCAAAACACGAGCAGAATGAAGCCTAGCC | Site-directed mutagenesis |
| ABD760 | MRI_T239E_F | CGCGGCTAGGCTTCATTCTGAGCGTGTTTTGGGAACATTC | Site-directed mutagenesis |
| ABD761 | MRI_T239E_R | GAATGTTCCCAAAACACGCTCAGAATGAAGCCTAGCCGCG | Site-directed mutagenesis |
| ABD800 | MRI_R240A_F | CTAGGCTTCATTCTACTGCTGTTTTGGGAACATTCGG | Site-directed mutagenesis |
| ABD801 | MRI_R240A_R | CCGAATGTTCCCAAAACAGCAGTAGAATGAAGCCTAG | Site-directed mutagenesis |
| ABD802 | MRI_R240S_F | CTAGGCTTCATTCTACTTCTGTTTTGGGAACATTCGG | Site-directed mutagenesis |
| ABD803 | MRI_R240S_R | CCGAATGTTCCCAAAACAGAAGTAGAATGAAGCCTAG | Site-directed mutagenesis |
| ABD804 | MRI_R240M_F | GCTAGGCTTCATTCTACTATGGTTTTGGGAACATTCGGC | Site-directed mutagenesis |
| ABD805 | MRI_R240M_R | GCCGAATGTTCCCAAAACCATAGTAGAATGAAGCCTAGC | Site-directed mutagenesis |
| ABD808 | MRI_K100E_F | GAAGCTGTTGCTATCGAAAAACTTGATGCTAG | Site-directed mutagenesis |
| ABD809 | MRI_K100E_R | CTAGCATCAAGTTTTTCGATAGCAACAGCTTC | Site-directed mutagenesis |
| JW17 | Mp_pEF1-F | GCAGTGGAGCGTCTGGCTTA | Genotyping of *proMp*EF1α-ORF fusions |
| JW25 | MpMRI_fwd_w/attB1 | GGGGACAAGTTTGTACAAAAAAGCAGGC TTAATGGCATGGTGTTGTTGCTG | Amplification of *Mp*MRI ORF with attB1-site |
| JW26 | MpMRI_rev_w/attB2 | GGGGACCACTTTGTACAAGAAAGCTGGG TTTCCATCCCGTGTGGGTGATC | Amplification of *Mp*MRI ORF with attB2-site and without stop-codon |
| JW34 | MpMRI_CDS_F1 | GCAACTCCAAGGCTGAGTGA | Genotyping of *Mp*MRI ORF |
| JW35 | MpMRI_CDS_R1 | CCTTGTCCACCTCCATGCAT | Genotyping of *Mp*MRI ORF |
| JW45 | Mp_rbm27_F1 | CCAAGTGCGGGCAGAATCAAGT | Genotyping of male Marchantia plants |
| JW46 | Mp_rbm27_R1 | TTCATCGCCCGCTATCACCTTC | Genotyping of male |

| | | | |
|---|---|---|---|
| | | | Marchantia plants |
| JW47 | Mp_rhf73_F1 | TGACGACGAAGATGTGGATGAC | Genotyping of female Marchantia plants |
| JW48 | Mp_rhf73_R1 | GAAACTTGGCCGTGTGACTGA | Genotyping of female Marchantia plants |
| JW52 | MpFER_3'UTR_R1 | CACTCCCAAATGAACGCACG | Genotyping of *Mpfer-1*/*Mp*FER |
| JW59 | MpFER_3'UTR_F2 | CGTGCCCTCTGTCTCCTGTTCC | Genotyping of *Mpfer-1*/*Mp*FER |
| JW60 | MpMRI_5'UTR_R1 | GCGGGCCTTGACTGCCTCT | Genotyping of *Mpmri-1/Mp*MRI |
| JW63 | pCambia1300_RB1 | CCTGCAGGCATGCAAGCTTGG | Genotyping of *Mpmri-1/Mp*MRI |
| JW64 | pCambia1300_RB2 | GCTGGCGTAATAGCGAAGAGG | Genotyping of *Mpmri-1/Mp*MRI |
